# Single-cell analysis of human retina identifies evolutionarily conserved and species-specific mechanisms controlling development

**DOI:** 10.1101/779694

**Authors:** Yufeng Lu, Fion Shiau, Wenyang Yi, Suying Lu, Qian Wu, Joel D. Pearson, Alyssa Kallman, Suijuan Zhong, Thanh Hoang, Zhentao Zuo, Fangqi Zhao, Mei Zhang, Nicole Tsai, Yan Zhuo, Sheng He, Jun Zhang, Genevieve L. Stein-O’Brien, Thomas D. Sherman, Xin Duan, Elana J. Fertig, Loyal A. Goff, Donald J. Zack, James T. Handa, Tian Xue, Rod Bremner, Seth Blackshaw, Xiaoqun Wang, Brian S. Clark

## Abstract

The development of single-cell RNA-Sequencing (scRNA-Seq) has allowed high resolution analysis of cell type diversity and transcriptional networks controlling cell fate specification. To identify the transcriptional networks governing human retinal development, we performed scRNA-Seq over retinal organoid and *in vivo* retinal development, across 20 timepoints. Using both pseudotemporal and cross-species analyses, we examined the conservation of gene expression across retinal progenitor maturation and specification of all seven major retinal cell types. Furthermore, we examined gene expression differences between developing macula and periphery and between two distinct populations of horizontal cells. We also identify both shared and species-specific patterns of gene expression during human and mouse retinal development. Finally, we identify an unexpected role for *ATOH7* expression in regulation of photoreceptor specification during late retinogenesis. These results provide a roadmap to future studies of human retinal development, and may help guide the design of cell-based therapies for treating retinal dystrophies.

## Introduction

The vertebrate retina is an accessible system for studying central nervous system (CNS) development. The retina develops from a polarized layer of neuroepithelial cells, that gives rise to six major classes of neurons and one class of glia in temporally distinct, but often overlapping, intervals during development. Certain cell types, such as retinal ganglion cells (RGCs), horizontal cells, cone photoreceptors, and GABAergic amacrine cells, are born relatively early, while glia, bipolar cells, glycinergic amacrine, and most rod photoreceptors are born relatively late (Cepko, 2014; La Vail et al., 1991; Voinescu et al., 2009; Wong and Rapaport, 2009; Young, 1985). The birth order of these cell types is evolutionarily conserved, and regulated by largely intrinsic mechanisms (Gomes et al., 2011; He et al., 2012).

Despite the high level of evolutionary conservation among many aspects of retinal development, there are important species-specific differences, particularly with respect to cell type composition and distribution. For instance, species differ substantially in the number of subtypes of these major cells, with mice having only one type of horizontal cell, but macaque and chicks having two or three, respectively (Boije et al., 2016). The relative ratio of rods to cones, and of photoreceptors to inner retinal cell types, also varies substantially (Peichl, 2005). Some species, such as mice, lack any major variation in the distribution of prominent cell types across the retina (Jeon et al., 1998) while others, such as humans and other primates, have a fovea in the central retina that is specialized for high-acuity vision and enriched in cones and retinal ganglion cells (Collin, 1999). The gene regulatory networks that control how these species-specific differences arise during development are still poorly understood.

An understanding of species-specific mechanisms controlling human retinal development is particularly important for therapies aimed at treating human-specific diseases. Two notable examples are retinoblastoma and macular degeneration. Retinoblastoma is caused by inactivation of the *RB1* tumor suppressor gene. The retina is exquisitely sensitive to *RB1* loss in humans, whereas it is highly resistant in other species examined to date. Murine models of retinoblastoma require the combined loss of *Rb1* and an additional tumor suppressor (Chen et al., 2004; Dannenberg et al., 2004; MacPherson et al., 2004; Sangwan et al., 2012; Zhang et al., 2004). Whereas the murine disease originates from ectopically dividing inner retinal neurons, human retinoblastoma has a cone origin (Xu et al., 2009, 2014). These phenotypic differences reflect disparities in the molecular circuitry of human versus murine cones (Xu et al., 2009). Since Rb1 inactivation promotes many tumors, a deeper understanding of human and murine cone circuitry could expose new insights into tumorigenesis. (Ajioka et al., 2007; Bremner and Sage, 2014; Chen et al., 2004; Xu et al., 2014)

Macular degeneration, which in its age-related (AMD) form affects up to 25% of the US population aged over 80 and is the most common form of age-related photoreceptor dystrophy (Jager et al., 2008), results in central vision loss from the selective loss of photoreceptors in the foveal region of the macula (Curcio et al., 2005). The relevance of animal models for AMD remains unclear since the macula is specific to primates, and most animal models lack regions of high-acuity vision. Although both bulk RNA-Seq analysis and small-scale single-cell RNA-Seq (scRNA-Seq) studies (Hoshino et al., 2017; Hu et al., 2019) have been used to profiled gene expression changes during retinal neurogenesis, these data have not shed light on human-specific mechanisms that regulate retinal development, particularly with respect to cone photoreceptor specification and foveal patterning.

The emergence of single-cell RNA sequencing (scRNA-Seq) technologies provides a powerful tool to comprehensively classify cell types of the central nervous system and the gene regulatory networks that control their development (Fan et al., 2018; Farrell et al., 2018; Liu et al., 2017; Tasic et al., 2018; Wagner et al., 2018; Zeisel et al., 2018; Zhong et al., 2018). Studies in both macaque and human have examined the diversity of cellular subtypes within the mature retina (Cowan et al., 2019; Lukowski et al., 2019; Peng et al., 2019; Voigt et al., 2019). Furthermore, recent studies in mouse using scRNA-Seq analysis have identified changes in gene expression across retinal development, including progenitor maturation and specification/differentiation of each of the major classes of retinal cell types (Buenaventura et al., 2019; Clark et al., 2019; Lo Giudice et al., 2019). These studies have led to the identification of genes such as the NFI family of transcription factors, which directly regulate retinal neurogenesis and cell fate specification. These large datasets also have the potential to identify both evolutionarily conserved and species-specific gene regulatory networks that control human retinal development.

Therefore, in this study, we employ a similar approach to generate a comprehensive scRNA-Seq profile of human retinal development. We profile 20 stages of human retinal organoid and *in vivo* development, ranging from early neurogenesis through adulthood, analyzing 118,555 cells retinal cells in total. Comparing human and mouse, we observe broadly similar changes in the gene expression profiles of both retinal progenitor cells (RPCs) and most postmitotic retinal cell types, but we also observe some major species-specific differences. These include the expression of transcription factors that control the specification of cone photoreceptors and horizontal interneurons, and also gene regulatory networks that pattern the fovea. Most notably, we find that the neurogenic bHLH factor *ATOH7*, which is selectively expressed in early-stage neurogenic RPCs and regulates the formation of early-born cells in many species including humans, is also expressed in late-stage neurogenic RPCs and regulates human photoreceptor specification. This resource and our analyses underscore the importance of obtaining gene expression data directly from primary human cells, illustrates the limitations of existing animal models for studying human disease, and provides insight into therapeutic approaches for treating retinal diseases.

## Results

### Construction and analysis of human retinal scRNA-Seq libraries

To comprehensively profile gene expression changes across retinal development, we performed scRNA-Seq on dissociated retinas from both human retinal organoids and primary tissues. To profile very early stages of retinal development, we profiled human retinal organoids, generated as previously described (Eldred et al., 2018), at 24, 30, 42 and 59 days *in vitro*. These timepoints correspond to early retinogenesis *in vivo* for which we were unable to procure samples, and include developmental windows of the earliest RPCs and initial RGC differentiation. Strikingly, a majority of the cells from organoids Days 24-42 in culture (13552/19861; 68.2%; Table S1) were annotated as non-eye-field cells due to expression of ventral forebrain and hypothalamic markers (*FOXG1, NKX2-1*, *DLX5, ARX, LHX6;* (Shimogori et al., 2010)) and lack of expression of eye-field specification markers (*VSX2, PAX6, RAX,* and *LHX2*). In addition, we generated profiles from whole developing retinas obtained at 9, 11, 12, 13, 14, 15, 16, 17, 18, 19, 22, 24, and 27 gestational weeks (GW), macular and peripheral samples from 20 gestational weeks and 8 days postnatal (PND), and whole retina from a healthy 86 year old donor (Fig. 1A; Fig S1A, Table S1). Samples were profiled to a mean depth of 3472.61 unique molecular identifiers (UMIs; standard deviation = 1862.78) and 1502.14 genes (standard deviation = 589.41) per cell (Fig. S1A-C). These samples were then integrated using Monocle 2.99.3 and UMAP dimension reduction on high variance genes to obtain a 3-dimensional embedding of retinal development (Fig. 1B,C; Fig. S1D,E; Movie S1-2; Table S2) (Becht et al., 2018; Qiu et al., 2017). Major cell types were annotated using previously identified marker genes (Blackshaw et al., 2004; Clark et al., 2019; Macosko et al., 2015; Siegert et al., 2012) expressed within clusters of cells in the dimension reduction space (Fig. S1F,G).

**Figure 1.**
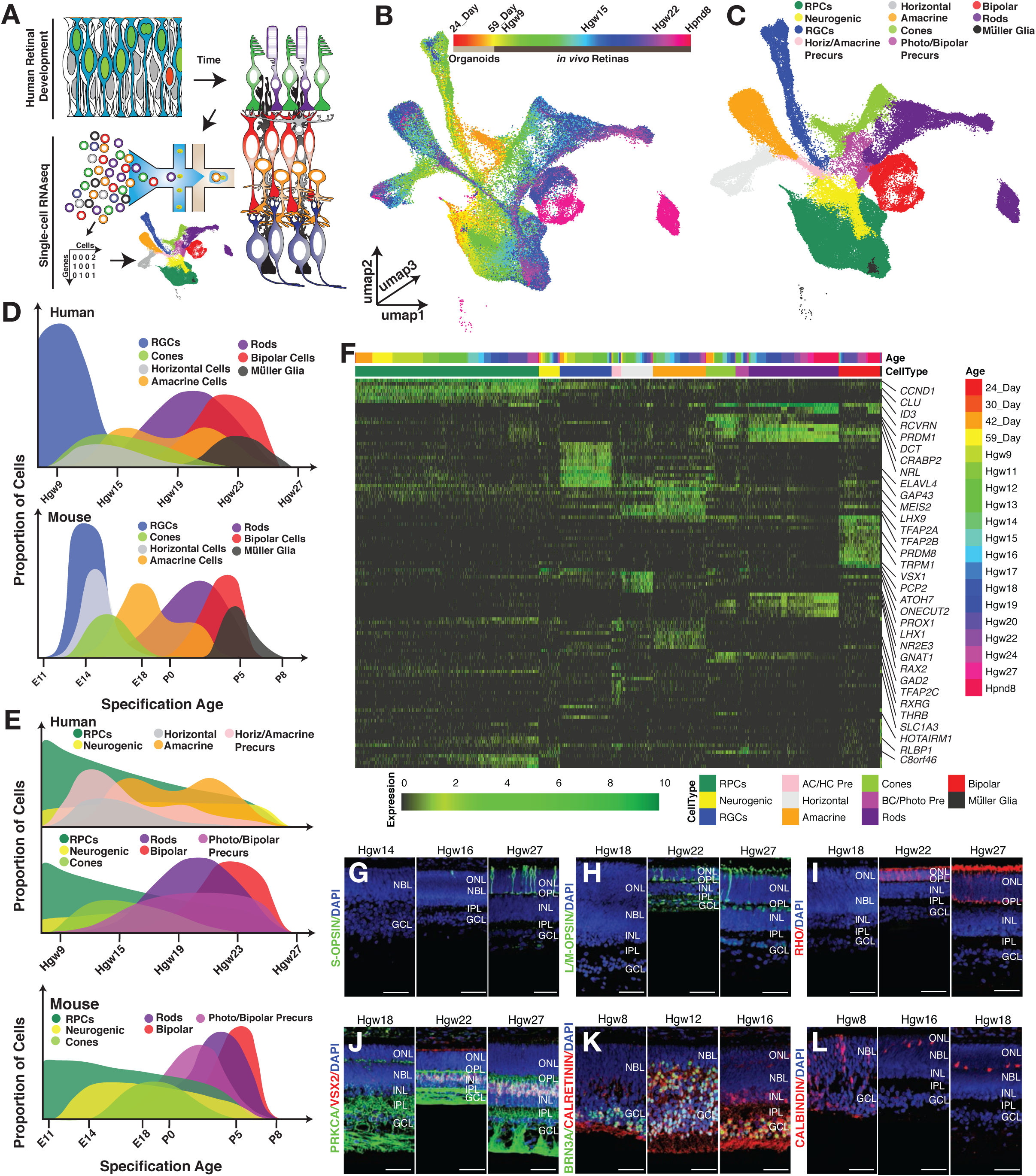
Single-cell RNAseq profiling of the developing human retina. (A) Schematic of experimental design. (B,C) 3D UMAP embedding of the retina dataset, with individual cells colored by (B) age and (C) annotated cell types. (D-E) Comparisons of cell type specification windows across human (top) and mouse (bottom) as inferred from the retinal scRNA-Seq datasets. (E) Precursor populations including neurogenic RPCs, amacrine/horizontal precursors and bipolar/photoreceptor precursor populations preceed peak generation of mature cell types. (F) Heatmap showing relative expression of transcripts with high specificity to individual cell types, ordered by cell type and developmental age (top annotation bars). (G-L) Immunohistochemistry on primary human retinal tissue validating the dynamic expression of cell-type markers, including (G) S-OPSIN (short wavelength cones); (H) L/M-OPSIN (long/medium wavelength cones); (I) RHO (rods); (J) PRKCA and VSX2 (bipolar cells); (K) BRN3A (RGCs) and Calretinin (Horizontal, Amacrine and RGC cells.); and (L) Calbindin (Horizontal cells). Nuclei are counterstained with DAPI. Scale bar: 50 μm. Abbreviations: Hgw - human gestational weeks; Hpnd - human postnatal day; RPCs - retinal progenitor cells; RGCs - retinal ganglion cells; AC/HC Pre - amacrine cell/horizontal cell precursors; BC/Photo Pre - bipolar cell/photoreceptor cell precursors; NBL - neuroblast layer; GCL - ganglion cell layer; ONL - outer nuclear layer; OPL - outer plexiform layer; INL - inner nuclear layer; IPL - inner plexiform layer.

Although these data were obtained from three different sources (Fig. S1A), involving different dissociation techniques, this nonetheless yielded an integrated cell distribution and set of developmental trajectories that broadly resemble those seen in mouse (Clark et al., 2019). Two major classes of RPCs are observed, which match the previously described primary and neurogenic subtypes in mouse (Clark et al., 2019). Differentiating Müller glia form a continuous developmental trajectory that emerges from the primary RPCs, while all retinal neurons emerge from the neurogenic fraction. As in the mouse, three major neuronal trajectories are observed: RGCs, amacrine and horizontal interneurons, and rods, cones and bipolar cells, respectively (Fig 1B-C; Fig S1D-E; Movie S2). RPCs and RGCs from organoids seamlessly integrate into this distribution, comprising the earliest developmental ages. Unlike in the mouse, we observed a distinct trajectory of human horizontal cells, likely due to the substantially increased abundance of captured horizontal cells within this dataset compared to mouse (5.8% in human versus 1.5% in mouse).

We next compared the fraction of each progenitor and postmitotic cell type observed at each time point to that in mouse (Fig. 1D,E; Fig. S1F-G) (Clark et al., 2019), and observed a broadly similar pattern of temporal windows for cell type specification. In both species, RGCs predominate early, with cones and horizontal cells being generated shortly thereafter. Peak relative numbers of rods and amacrine cell generation occurred near the midpoint of neurogenesis, while bipolar and Müller glial cells emerged last (Fig. 1D). In humans, however, a broader temporal distribution of cone and horizontal cell generation was observed. Additionally, we examined the ages at which primary RPC, neurogenic RPC, and cell type precursors are observed across the human and mouse scRNA-Seq datasets, highlighting temporal windows of amacrine, horizontal, bipolar, and photoreceptor cell specification windows (Fig. 1E).

We next identified selective markers of each major progenitor and postmitotic cell type by feature ranking in clustered single cell data using genesorteR (Ibrahim and Kramann, 2019) (Fig. 1F). This revealed known and novel markers of specific cell types. While most show identical cell-type specificity in both mice and humans, additional or entirely novel patterns of cell-specific expression were evident in some cases. For instance, in mice, the lipid-binding protein *CLU* is selectively expressed in Müller glia (Blackshaw et al., 2004), but in humans it is strongly expressed in both primary RPCs and Müller glia. In contrast, while retinoid binding protein *CRABP2* is expressed exclusively in early-stage primary RPCs in mouse (Clark et al., 2019), it is also strongly expressed in human cone precursors. Other examples of cone-enriched genes in humans that show different patterns of expression in mouse include *DCT*, which is restricted to the retinal pigmented epithelium in mouse, and *HOTAIRM1*, which is not detectably expressed in mouse retina (Fig. 1F). Other differences include human-specific genes, such as *RAX2,* which we detected in both cone and rod photoreceptors.

Immunohistochemical analysis demonstrates that protein expression patterns of known marker genes also broadly reflect cell-specific and temporally dynamic patterns of transcript expression detected by scRNA-Seq (Fig. 1G-L; Fig. S1H-N).

### Early and late-stage human RPCs show distinct gene expression profiles

Previous scRNA-Seq analysis of mouse retinal development identified clear transcriptional signatures of early and late-stage primary RPCs (Clark et al., 2019), reflecting differential expression of genes that control proliferation, neurogenesis, and cell fate determination. To determine the extent of evolutionary conservation in temporal patterning of RPCs, we conducted pseudotemporal analysis of gene expression changes across primary RPCs and Müller glia. Interestingly, organoid-derived RPCs displayed gene expression patterns of early-stage RPCs from primary tissue (Fig. 2A-C). Pseudotemporal ordering of RPCs identified bimodal densities of RPCs across pseudotime, followed by Müller glia that reflected the sequential developmental ages of input tissue (Fig. 2A-C; S2A,B). This mirrored changes observed in mouse, although the transition between early and late-stage RPCs occurred more gradually. In mice, the transition between early and late-stage RPCs occurs rapidly between E16 and E18, but in humans this process occurs between 11 and 15 gestational weeks (GW), and likely reflects the regional differences in developmental human retina development between central and peripheral regions (Diaz-Araya and Provis, 1992; van Driel et al., 1990). The fraction of human RPCs in G1 is increased relative to mouse (*p* < 0.0001; *chi*-squared test; Fig. S2C), which may reflect the much longer time course of retinal neurogenesis seen in humans relative to mice (Centanin and Wittbrodt, 2014).

**Figure 2.**
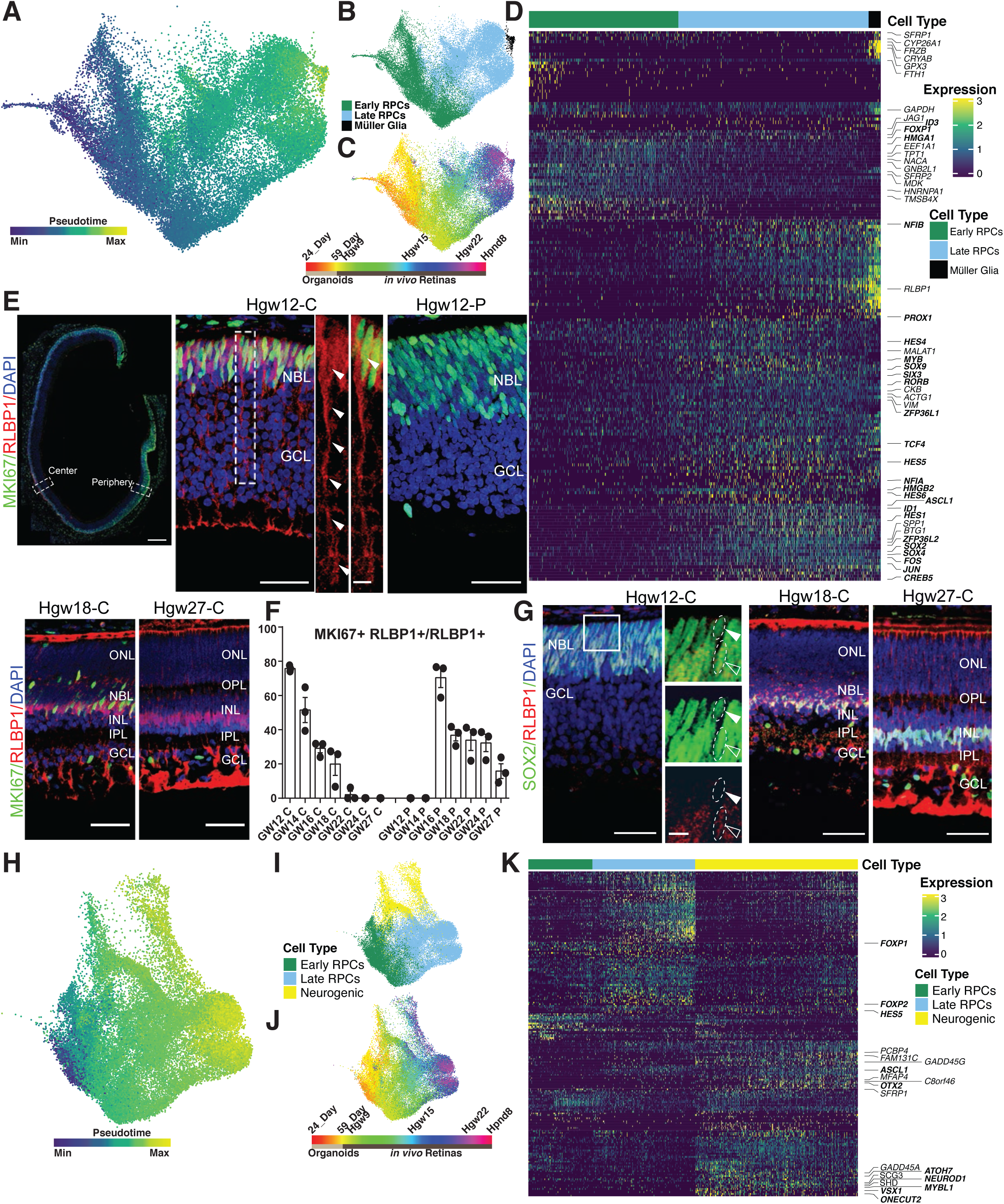
Pseudotime analysis reveals genes differentially expressed across primary and neurogenic RPCs. (A-C) UMAP embedding of the developmental trajectories of primary RPCs and Müller glia with cells colored by (A) pseudotime, (B) cell type and (C) developmental age. (D) Heatmap of differentially expressed transcripts along pseudotime from primary RPC to Müller glia. Cells are ordered by cell type and pseudotime with transcription factors listed in bold. (E) Immunohistochemistry detecting for RLBP1 and MKI67 in Hgw12 (top panels), Hgw18-Central (bottom left), and Hgw27-Central retina (bottom right) with magnified views of central (top center) and peripheral (top right) regions. Nuclei are counterstained with DAPI. Hgw12 scale bar: 300μm (left), 50μm (center and middle), 10μm (center side panels); Hgw18-C and Hgw27-C scale bar: 50μm. (F) Bar chart showing the proportion of actively proliferating (MKI67+) cells amongst the RLBP1+ population in the central and peripheral regions across the developing human retina (Hgw12, Hgw14, Hgw16, Hgw18, Hgw22, Hgw24, and Hgw27). Data are mean ± SEM. (G) Immunohistochemistry of SOX2 and RLBP1 in central regions of developing human retinas at Hgw12, 18 and 27. High magnification images are of the boxed region in the Hgw12 image. Open arrowheads indicate co-localization, with closed arrowheads indicating failure to detect *RLBP1* expression. Nuclei are counterstained with DAPI. Hgw12 scale bar: 50μm (left), 10μm (right); Hgw18-C and Hgw27-C scale bar: 50μm.(H-J) UMAP embedding of primary and neurogenic RPCs with cells colored by (H) pseudotime, (I) cell type and (J) developmental age. (K) Heatmap of differential transcript expression along pseudotime from primary RPC to neurogenic RPC. Cells are ordered by cell type and pseudotime with transcription factors listed in bold. Abbreviations: Hgw - human gestational weeks; GW - gestational weeks; Hpnd - human postnatal day; NBL - neuroblast layer; GCL - ganglion cell layer; ONL - outer nuclear layer; OPL - outer plexiform layer; INL - inner nuclear layer; IPL - inner plexiform layer; C - central retina; P - peripheral retina.

We observed similar expression of markers for early and late-stage primary RPCs in human as in mouse, including *SFRP2* and *NFIA* (Fig. 2D; Fig. S2G). Likewise, many of the genes upregulated in Müller glia begin their expression in late-stage primary RPCs (i.e *RLBP1*, *NFIB*). Unlike the Müller glia-specific expression pattern seen in mice (Blackshaw et al., 2004; Reichenbach and Bringmann, 2013), however, both *CLU* and *VIM* are expressed in primary RPCs at all stages of neurogenesis (Fig. 1F; Fig. 2D). Known inhibitors of the WNT pathway (*SFRP1* and *FRZB*) are expressed in early RPCs, and then again specifically in Müller glia at later stages. Immunohistochemistry revealed that expression of RLBP1 is observed in KI67^+^ mitotic late-stage RPCs in central retina by 12 GW and displays a more developmentally delayed peripheral retinal expression by 16 GW (Fig. 2E,F; Fig. S2D). The loss of MKI67 and RLBP1 co-localization coincided with the emergence of co-localization of SOX2 and RLBP1 in Müller glia (Fig. 2G; Fig. S2E-F).

Neurogenic RPCs are observed in all samples except the Day24 organoids and PND8 samples (Fig. 2H-J, Tables S1), and pseudotemporal ordering of neurogenic RPCs reflected transcriptional signatures of age-matched primary RPCs. Neurogenic RPCs likewise showed broadly similar temporal expression patterns in humans as in mice (Fig. S2H). As in mouse, a subset of human genes show selective expression in either early or late-stage neurogenic RPCs (Fig. 2J,K; Fig. S2H). However, pseudotime analyses did not reveal a clear signature of early and late-stage human neurogenic RPCs as observed in mouse (Fig. S2J-K; (Clark et al., 2019)). Genes that display temporal enrichment in early-stage neurogenic RPCs in both humans and mice include *DLX1/2, GAL*, *ONECUT2*, and *ATOH7*, while genes enriched in late-stage neurogenic RPCs include *ASCL1*, *OTX2*, and *SOX4* (Fig. 2K; Fig. S2H). A handful of genes, however, showed substantial species-specific differences in expression. Several genes that are expressed across neurogenic RPC development in mouse -- such as *OLIG2*, *NEUROG2*, and *BTG2* -- display higher expression in late-stage human neurogenic RPCs (Fig. S2I). Moreover, *C8ORF46* (*3110035E14Rik*; *Vexin*), which is selectively expressed in early-stage neurogenic RPCs in mouse (Clark et al., 2019; Moore et al., 2018), is also expressed in late-stage neurogenic RPCs in humans (Fig. S2I). A small number of genes that lack mouse orthologs, including *HES4*, are also enriched in human neurogenic RPCs (Fig. S2I). As in primary RPCs, a greater fraction of human neurogenic RPCs are in G1 than in mouse (p < 0.0001; Chi-square test), and the fraction of cells in G1 increases steadily over the course of neurogenesis (Fig. S2L).

### Identification of differentiation trajectories for each major human retinal cell type

To identify gene regulatory networks that control retinal cell specification and differentiation, we independently conducted pseudotime analysis along the cellular trajectories of each major retinal cell type (Fig. 3; Fig. S3). We observed many transcription factors that show highly-specific expression at various stages of differentiation of each cell type in humans, display similar expression in mouse. Both human and mouse retinal ganglion cells express the well-characterized transcript markers *POU4F1, POU4F2, POU6F2, ISL1, NHLH2, RXRG, EBF1* and *EBF3* (Clark et al., 2019; Rheaume et al., 2018; Tran et al., 2019); Fig. 3A-D); however, human RGCs express MYC (Fig. 3A). Human horizontal cells express *TFAP2A/B*, *ONECUT2/3*, *LHX1* and *ESRRB*, as previously observed in mouse (Fig. 3E-H) (Boije et al., 2016; Clark et al., 2019), and show considerable overlap in transcription factor expression with amacrine cells (Fig. 3I-L). However, differentiating human starburst amacrine cells (SACs) selectively express *NEUROG3*, which is not expressed in mouse retina in any cell type, but human SACs also express canonical transcription factors *SOX2, ISL1,* and *FEZF1* (Fig. S3A-C) (Balasubramanian and Gan, 2014; Clark et al., 2019).

**Figure 3.**
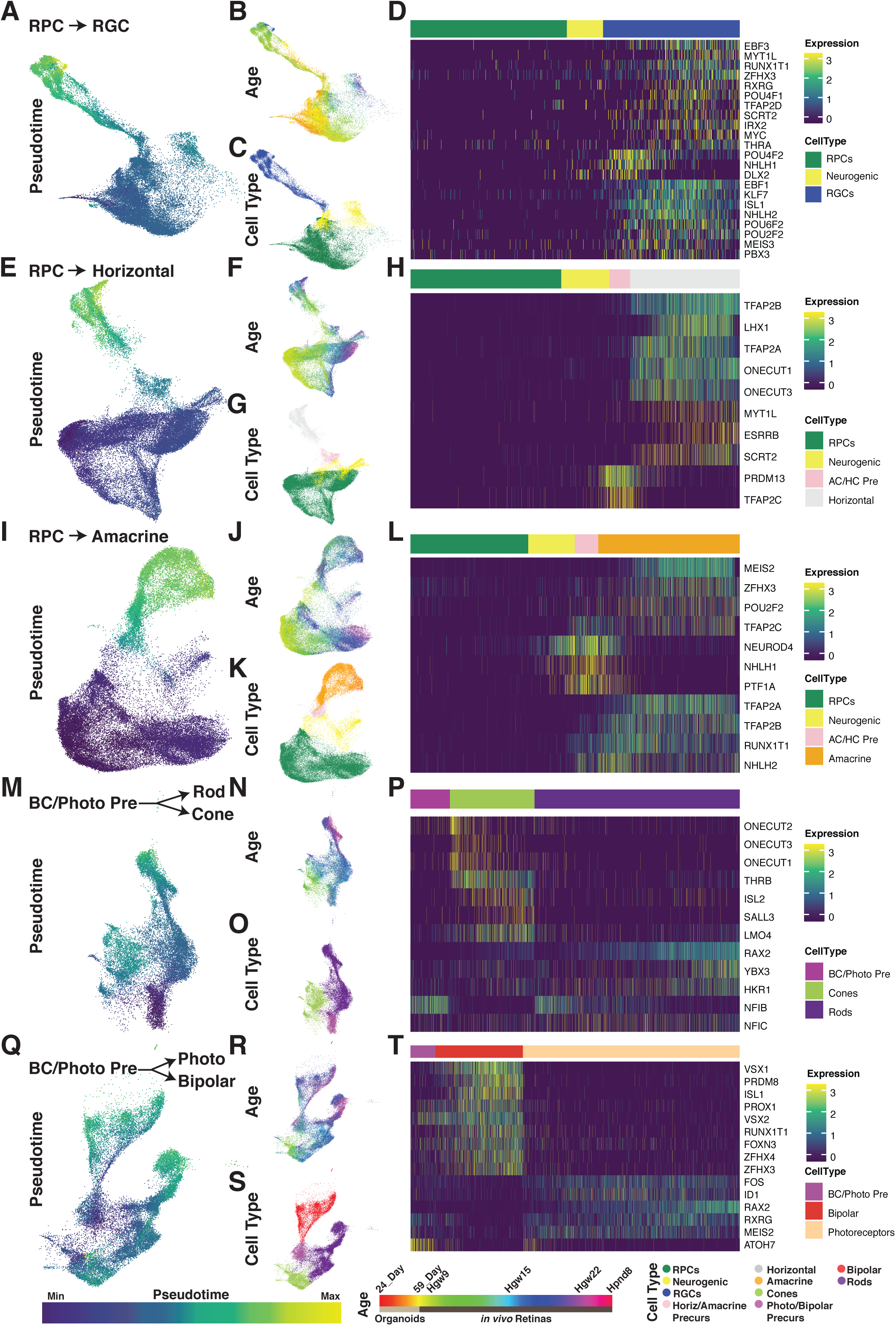
Pseudotime analyses identify transcription factor networks controlling human retinal cell fate specification. UMAP embeddings of cellular inputs for pseudotime analyses of (A-D) retinal ganglion cells, (E-H) horizontal cells, (I-L) amacrine cells, (M-P) rods/cones, and (Q-T) photoreceptor/bipolar cells. UMAP plots are colored by (A,E,I,M,Q) cellular pseudotime values, (B,F,J,N,R) age, and (C,G,K,O,S) cell type. (D, H, L, P, T) Heatmap showing relative expression of differentially expressed transcription factors across pseudotime, highlighting transcription factors with enriched expression in endpoint cell types. Abbreviations: Hgw - human gestational weeks; Hpnd - human postnatal day; RPC - retinal progenitor cells; RGC - retinal ganglion cells; Photo - photoreceptors; BC/Photo Pre - bipolar cell/photoreceptor precursors.

Some of the more pronounced differences in transcription factor expression between human and mouse are observed in differentiating photoreceptors. In the mouse, *ISL2* is strongly expressed in retinal ganglion cells, but only weakly expressed in cone photoreceptors (Triplett et al., 2014). In both humans and chick, however, (Edqvist et al., 2006; Triplett et al., 2014), *ISL2* is strongly expressed in human cones, along with its cofactor *LMO4* (Fig. 3M-P). The Kruppel class zinc finger transcription factor *HKR1*, which is selectively expressed in developing human rods, is entirely absent from the mouse genome. Strikingly, *ATOH7* is also expressed in human photoreceptor/bipolar precursors and immature cones (Fig. 3Q-T), in sharp contrast to the mouse and also every other species examined to date, where it is preferentially expressed in early-stage neurogenic RPCs (Brown et al., 1998; Kanekar et al., 1997; Kay et al., 2001). These findings imply the existence of species-specific differences in the transcriptional regulatory networks controlling rod and cone photoreceptor specification.

Since we observed a organoid-specific cone trajectory in our dimension reductions (Fig. 1B-C), we next performed the same pseudotemporal analysis on organoid versus *in vivo* cone development,. This analysis of gene expression identifies some unexpected differences in the differentiation of cone photoreceptors in retinal organoids relative to primary tissue (Fig. S3D-F). While organoid derived cones express substantially lower levels of cone precursor-enriched transcription factors such as *CRX*, the organoid-derived cones also express neurogenic factors typically associated with non-photoreceptor cell types including *LHX9*, *NHLH1*, and *SOX11*. Subsets of organoid-derived cones, however, do express additional cone photoreceptor-enriched genes. These include *THRB, ISL2, LMO4, RXRG, SALL3* and *DCT*. Therefore, as indicated by the relative position of the organoid-cone trajectory within the dimension reduction, the organoid-derived cones more closely resemble native cones than any other cell type, (Fig. 1B-C; Movie S1-2).

While we identified many transcripts/transcription factors as differentially expressed across the pseudotemporal analyses of individual cell trajectories (Fig. 3, Fig. S3), it should be noted that many of the genes are not specific to any one cell trajectory (Fig. S3G), suggesting that many transcription factors regulate specification of multiple cell types.

### Central vs. peripheral differences in retinal transcript expression

To identify regional differences in gene expression associated with the development of the macula compared to the peripheral retina, we examined differential gene expression between macula and peripheral cells from the GW20 and PND8 retinal samples using the ‘fit_models’ function from Monocle3. We observed broad transcriptional differences between the regions within individual cell types (Fig. S4 A-I); however, analyses are partially confounded by cell type capture efficiencies inherent to the datasets. Furthermore, since macular retina is developmentally advanced relative to peripheral retina (Diaz-Araya and Provis, 1992; van Driel et al., 1990; La Vail et al., 1991), we examined the correlation of differentially expressed transcripts with pseudotime trajectories across each individual cell type (Fig. S4 J-Q). The correlation with pseudotime helps to distinguish bona fide enrichment in cells in macular retina from simple differences in developmental age between macular and peripheral retina. The pseudotime analyses suggest many of the observed regional differences are correlated (both positively and negatively) with cell type differentiation and maturation. However, other transcripts, such as the cone macula-enriched transcript *DST* display little correlation with pseudotime, and may reflect true regional differences.

We identified multiple genes that were differentially expressed between macular and peripheral RPCs that showed little correlation with pseudotime (correlation close to 0; Fig S4Q). These include *CYP26A1, DIO2, CDKN1A, ANXA2,* and *FRZB* (Fig. 4A; Fig. S4J-Q). Of particular interest is *CYP26A1,* a transcript implicated in marking a rod-free zone in the chicken (da Silva and Cepko, 2017), and selectively expressed in RPCs and Müller glia of both the developing and mature primate fovea (Cowan et al., 2019; Peng et al., 2019). We then validated enrichment of *CYP26A1* and *DIO2* transcript expressions and co-expression of *SFRP1* or *RLBP1* within macula cells of the GW18 retina (Fig 4B-C). This allowed us to extrapolate the expression of any of the 5 listed marker genes to the rest of the RPCs within the dataset to identify additional potential macular RPCs within whole retina dissociations (Fig. 4D). Subsequent differential tests were performed on the inferred macular/peripheral primary RPCs using a regression model to reduce the effect of cell cycle phase on differential gene expression. Our results highlight additional potential candidate regulators of macula development. Sample and cell type specific enrichment of the candidate regulators is shown in Figures 4E-F, many of which are enriched in RPCs and Müller glia. One interesting candidate macula development regulator gene enriched in our macular RPC population is *CTGF*. *CTGF is* a downstream target of the Hippo signaling pathway. While *CTGF* displays enriched expression within Müller glia, recent studies have identified a FGF15 (19 in humans)/FGFR4-mediated pathway for Hippo-pathway activation (Ji et al., 2019). Since *FGF19* expression marks early RPCs, activation of Hippo-signaling and degradation of retinoic acid (*CYP26A1* expression) may function concordantly to confer macular specialization. Macula-enriched expression is observed for genes expressed in all major cell types (Fig. 4F; Fig. S4J-Q). Comparison of genes enriched in developing human macula with foveal-enriched genes from adult macaque retina reveals that *CYP26A1* and *CTGF* are enriched in Müller glia in both species, as are *RBP3* and *PROM1* in cones, and *VTN* in rods (Peng et al., 2019).

**Figure 4.**
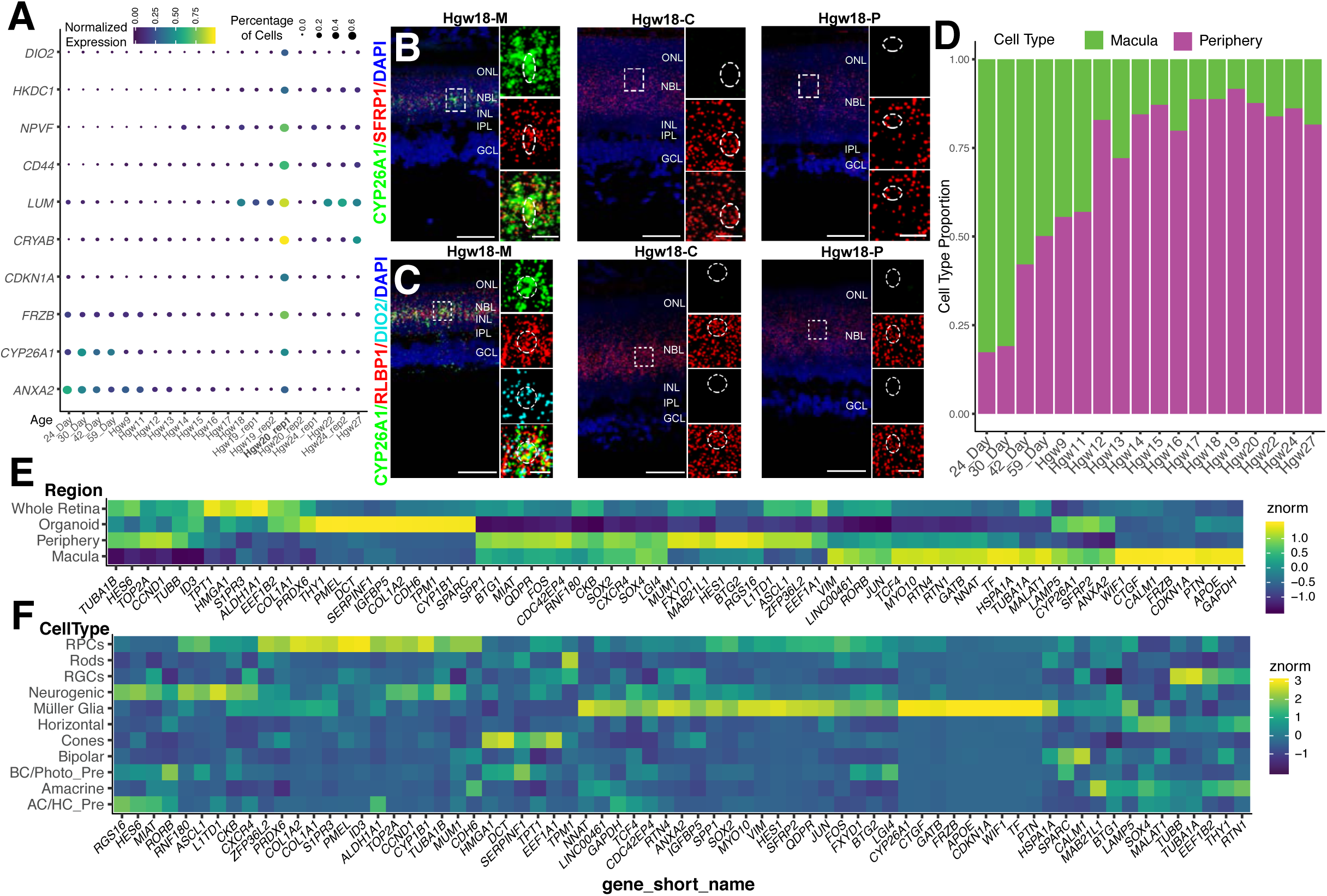
Identification of macular RPC transcripts for regional specification of the developing human retina. (A) Dotplot of differentially expressed genes between macular and peripheral retina RPCs and their relative expression and percentage of expressing cells in RPCs in each sample. The bolded Hgw20_rep1 sample highlights a macular sample containing significant numbers of RPCs. (B-C) RNAscope detecting (B) *CYP26A1* and *SFRP1* and (C) *CYP26A1*, *RLBP1* and *DIO2* transcripts in macular, central, and peripheral Hgw18 retina samples with high magnification images of boxed regions. Nuclei are counterstained with DAPI. Scale bar: 50μm and 10μm (magnified views). (D) Proportion of macular and peripheral RPCs as classified by *CYP26A1, CDKN1A, DIO2*, *ANXA2,* or *FRZB* expression at each age. (E-F) Heatmap showing (E) regional and (F) cell type expression enrichment of differentially expressed transcripts between the macula and peripheral RPCs. Abbreviations: Hgw - human gestational weeks; M - macular retina; C - central retina; P - peripheral retina; NBL - neuroblast layer; GCL - ganglion cell layer; ONL - outer nuclear layer; OPL - outer plexiform layer; INL - inner nuclear layer; IPL - inner plexiform layer; RPCs - retinal progenitor cells; AC/HC Pre - amacrine cell/horizontal cell precursors; BC/Photo Pre - bipolar cell/photoreceptor precursors, RGCs - retinal ganglion cells.

### Specification and differentiation of two human horizontal cell subtypes

While mouse retinas contain a single subtype of horizontal cell, primate retinas contain at least two distinct subtypes. Recent scRNA-Seq analysis of macaque retina identified two horizontal cell subtypes distinguished by the presence or absence of *CALB1* expression (Peng et al., 2019). Our analysis of human horizontal cell precursors also clearly identifies two subtypes of differentiating horizontal cells, distinguished by differential expression of the LIM homeodomain transcription factors *LHX1* and *ISL1* (Fig. 5A-C; Fig. S5F). Using smfISH, we confirm that *LHX1*+ and *ISL1*+ horizontal cells are distinct populations (Fig. 5D-H), but that both express the pan-horizontal cell markers *ONECUT1-3* (Fig. S5A-C; G,H). The presence of *ISL1*+ horizontal cells in humans is similar to the chicken, where 2 populations of Isl+ horizontal cells with distinct morphology are observed (Suga et al., 2009). The *ISL1*+ horizontal cells are also CALB1+, likely corresponding to the previously identified *CALB1*+ horizontal cells of macaque (Fig. S5D) (Peng et al., 2019). However, since all horizontal cells are both *Lhx1*+ and *Calb1*+ in mouse (Boije et al., 2016; Clark et al., 2019; Lo Giudice et al., 2019), this represents a major species-specific difference in gene expression. Differential expression and pseudotemporal analyses identify additional markers of these horizontal subtypes, including *PCDH9, C1QL1,* and *SOSTDC1* for *LHX1+* cells and *FAM135A* and *SYNPR* for *ISL+* horizontal cells (Fig. 5I; Fig. S5E,I). The two populations of horizontal cells also show different expression of cell surface receptors. *LHX1+* cells specifically expresses the dopamine receptor *DRD2,* while no other dopamine receptor genes are expressed in either horizontal cell population (Fig. 5I). *LHX1+* cells also express *MEGF11*, which has been shown to regulate mosaicism of horizontal cells in mouse (Kay et al., 2012), suggesting that an alternative mechanism is likely used to form ISL1+ horizontal cell mosaics (Fig. 5I).

**Figure 5.**
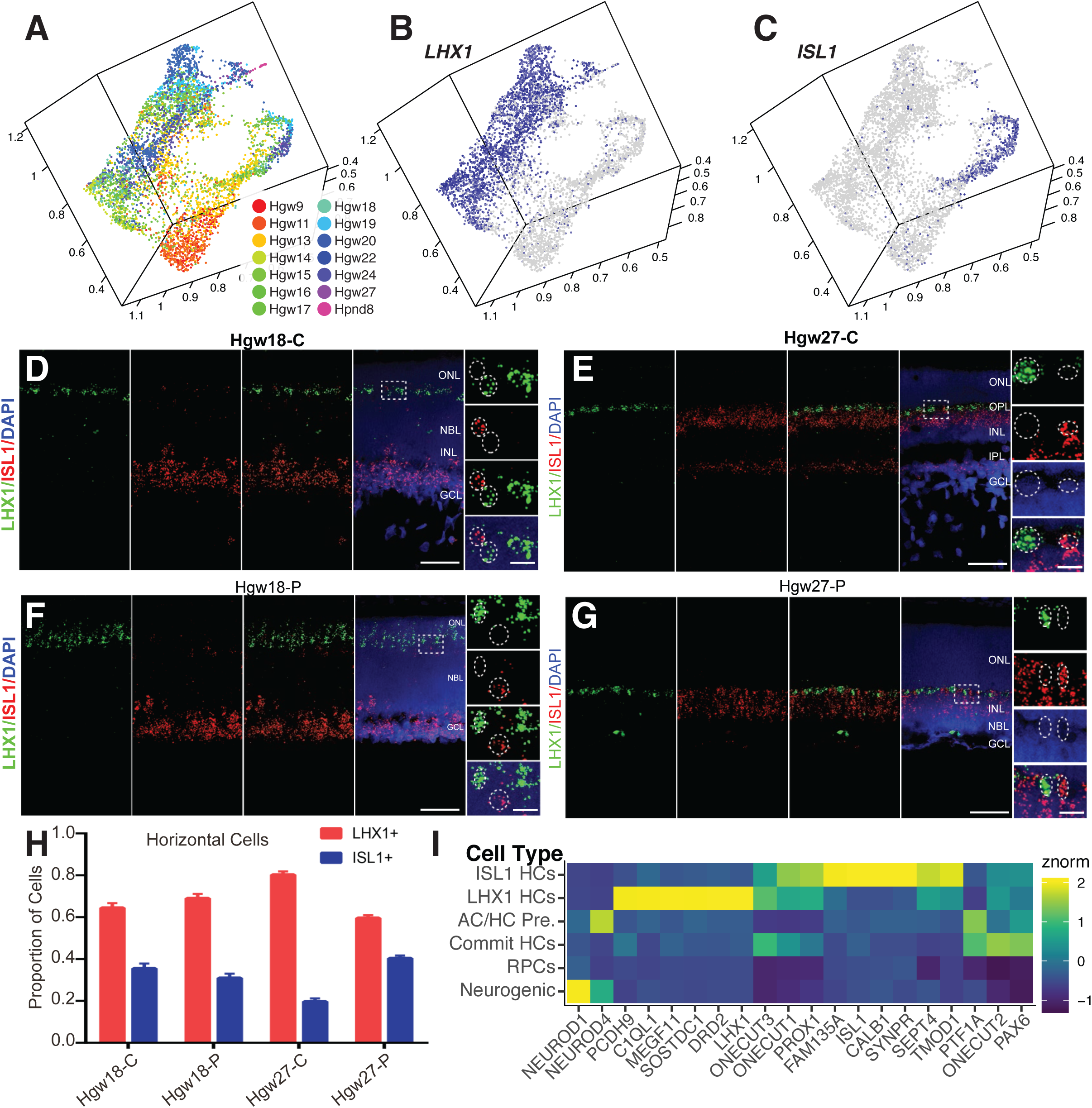
Identification and differentiation of two horizontal cell subtypes within the developing human retina. (A-C) UMAP embedding of Horizontal Cells, colored by (A) age or (B-C) relative expression of (B) *LHX1* and (C) *ISL1*. (D-G) RNAScope detecting expression of *LHX1* and *ISL1* transcripts in central (D,E) and peripheral (F,G) Hgw18 and Hgw27 human retina, with higher-magnification views of the boxed regions. Nuclei are counterstained with DAPI. Scale bars: 50 μm and 10 μm (magnified views). (H) Quantification of the proportions of each horizontal cell subtype in central and peripheral retina at ages Hgw18 and 27 from fluorescent *in situ* hybridization experiments. Data are mean ± SEM. (I) Heatmap showing relative cell type expression of horizontal cell commitment, differentiation and subtype specification genes. Abbreviations: Hgw - human gestational weeks; Hpnd - human postnatal day; C - central retina; P - peripheral retina; NBL - neuroblast layer; GCL - ganglion cell layer; ONL - outer nuclear layer; OPL - outer plexiform layer; INL - inner nuclear layer; IPL - inner plexiform layer; AC/HC Pre. - amacrine cell/horizontal cell precursors; Commit HCs - committed horizontal cells, ISL1 HCs - *ISL1*-positive horizontal cells; LHX1 HCs - *LHX1-*positive horizontal cells.

### Identification of convergent and divergent gene expression patterns in mouse and human retina using scCoGAPS and projectR

To identify gene regulatory networks in human retinal development, we used scCoGAPS on 3113 transcripts displaying high variance across the dataset (Table S2), identifying 112 patterns of gene co-expression in our human scRNA-Seq data (Sherman et al., 2019; Stein-O’Brien et al., 2019). Through projectR, we then projected our previously published mouse retinal development scRNA-Seq dataset, excluding all non-neuroretinal cells, onto these 112 human retinal patterns (Clark et al., 2019; Stein-O’Brien et al., 2019). By analyzing associations between each pattern and each main retinal cell types in both human and mouse datasets, we were able to identify patterns that are divergent or convergent between the two species (Fig. 6A; Fig. S6 A-D). In general, most patterns highly correlated with individual cell types showed broadly similar cellular expression patterns in both species. However, several patterns stood out as highly divergent, particularly patterns correlated with human cone development. One example of divergent patterns between mouse and human is Human Pattern 75, which marks human cones but highlights both early RPCs and cones in mouse (Fig. 6A-D; Fig. S6A,C). Examination of the top marker genes of human Pattern 75 indicate that the divergence between the species is driven by transcripts that include *CRABP2, DCT, and LOXL1* (Fig. 6E). Specifically, human *LOXL1* is expressed in developing neurons, with high expression in cones and developing rods (Fig. 6F). In contrast, mouse *Loxl1* shows little expression within the developing retina, being detected in only a few early-stage RPCs (Fig. 6G).

**Figure 6.**
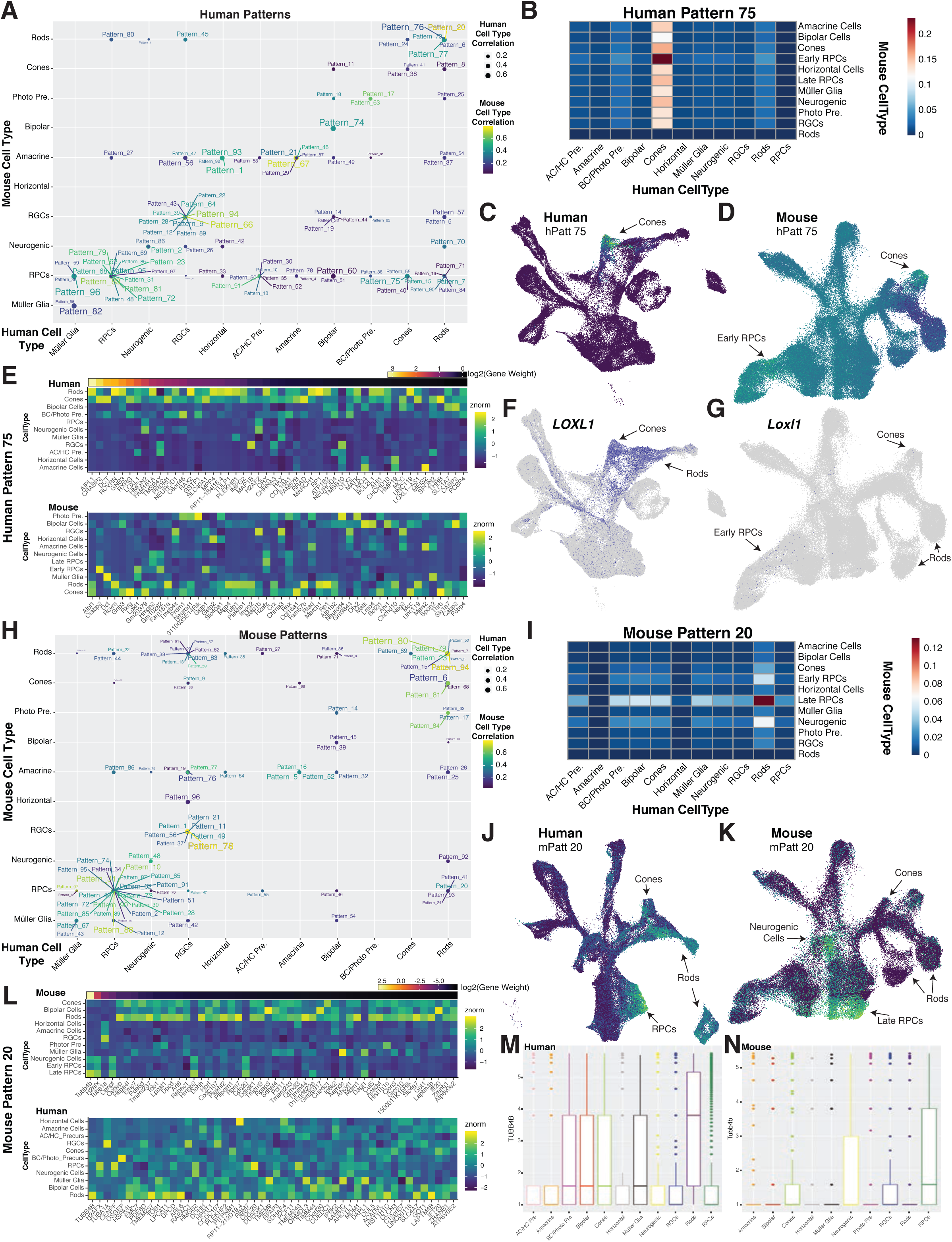
Cross-species comparisons of gene usage using scCoGAPS and projectR reveals conserved divergent pattern usage across human and mouse retinal cell types. (A) Plot indicating maximally correlated retinal cell types in mouse (y-axis) and human (x-axis) of human patterns. Correlation values are indicated through size of the point for human cell types, and color of the dot and label for mouse cell types. (B) Heatmap indicating the intersection of correlations of Human Pattern 75 for human (x-axis) and mouse (y-axis) cell types. (C-D) UMAP embeddings of (C) human and (D) mouse scRNA-Seq datasets, with cells colored by Human Pattern 75 pattern weights. (E) Heatmap of relative gene expression within (top) or (bottom) mouse cell types for the top 50 weighted genes of Human Pattern 75 and their orthologs in mouse (bottom). Genes are ordered by relative gene weights. (F-G) UMAP embeddings displaying *LOXL1*/*Loxl1* expression within (F) human and (G) mouse retinal scRNA-Seq datasets. (H) Plot of maximally correlated retinal cell types in mouse (y-axis) and human (x-axis) of mouse patterns. Correlation values are indicated through size of the point for human cell types, and color of the dot and label for mouse cell types. (I) Heatmap indicating the intersection of correlations of Mouse Pattern 20 for human (x-axis) and mouse (y-axis) cell types. (J-K) UMAP embeddings of (J) human and (K) mouse scRNA-Seq datasets, with cells colored by Mouse Pattern 20 pattern weights. (L) Heatmap of relative gene expression within (top) mouse or (bottom) human cell types for the top 50 weighted genes of Mouse Pattern 20 and their orthologs in human (bottom). Genes are ordered by relative gene weights. (M-N) Box-plots displaying the log2(expression + 1) of *TUBB4B* in each (M) human and (N) mouse retinal cell type. Abbreviations: AC/HC Pre. - amacrine cell/horizontal cell precursors; RGCs - retinal ganglion cells; RPCs - retinal progenitor cells; BC/Photo Pre - bipolar cell/photoreceptor precursors; Photo Pre - photoreceptor precursors; hPatt - human pattern; mPatt - mouse pattern.

We also identified 97 gene patterns across 3164 highly variable genes within the neuroretinal cells from the mouse scRNA-Seq dataset (Table S2) and repeated the same comparison process (Clark et al., 2019; Stein-O’Brien et al., 2019). Of note, 753 of the >3000 highly variable were used as input to scCoGAPS across both species. A similar overall picture was seen in mouse, with patterns displaying high correlation with individual cell types corresponding to the same cell type in humans. Some patterns displayed low correlation with cell types across the species (Fig. 6H; Fig. S6E-H). Notable among these is Pattern 20, which in mouse is enriched in late-stage RPCs but in human is enriched in rods, cones and late-stage RPCs (Fig. 6H-K; Fig. S6E,G). *Tubb4b*, a marker for Pattern 20, shows substantial differences in cellular expression levels in late-stage RPCs, rods and cones in humans and mice (Fig. 6L-N). These unbiased cross-species analyses highlight both similarities and disparities in gene usage across mouse and human retinal development, in particular in the control of photoreceptor specification and differentiation, and can be used to identify instances where mouse models may not recapitulate human disease.

### ATOH7 controls the relative ratio of human rod and cone photoreceptors

Multiple lines of evidence from this study suggest that the gene regulatory networks controlling photoreceptor development differ substantially between mouse and humans. We identified *ATOH7* expression in late-stage neurogenic RPCs and immature cones, in addition to its expression in early-stage neurogenic RPCs, which has been observed in every other species examined to date (Fig. 7A; Fig. S7A,B) (Brown et al., 1998; Kanekar et al., 1997; Kay et al., 2001; Matter-Sadzinski et al., 2001). This was confirmed using immunostaining for ATOH7 and the early photoreceptor-enriched markers OTX2 and CRX. At both early and late stages of photoreceptor development, we observe fraction of both OTX2 (Fig. 7B,C; Fig. S7E) and CRX-positive cells (Fig. 7D,E; Fig. S7F) that also express ATOH7. This colocalization was temporally dynamic and showed major differences between central and peripheral retina. Nearly 10% of all OTX2+ cells in central retina were also ATOH7-positive at GW10, but at later stages very little colocalization was observed (Fig. 7C). In contrast, at stages examined between GW10 and GW20, 12-20% of all OTX2-positive cells in peripheral retina were also ATOH7-positive. Colocalization of ATOH7 and CRX also showed a large difference between central and peripheral retina, with very little observed in central retina, but more than 30% of CRX-positive cells in peripheral retina expressing ATOH7 at GW14, though with the fraction of co-expressing cells declining rapidly thereafter to undetectable levels at GW22 (Fig. 7E).

**Figure 7.**
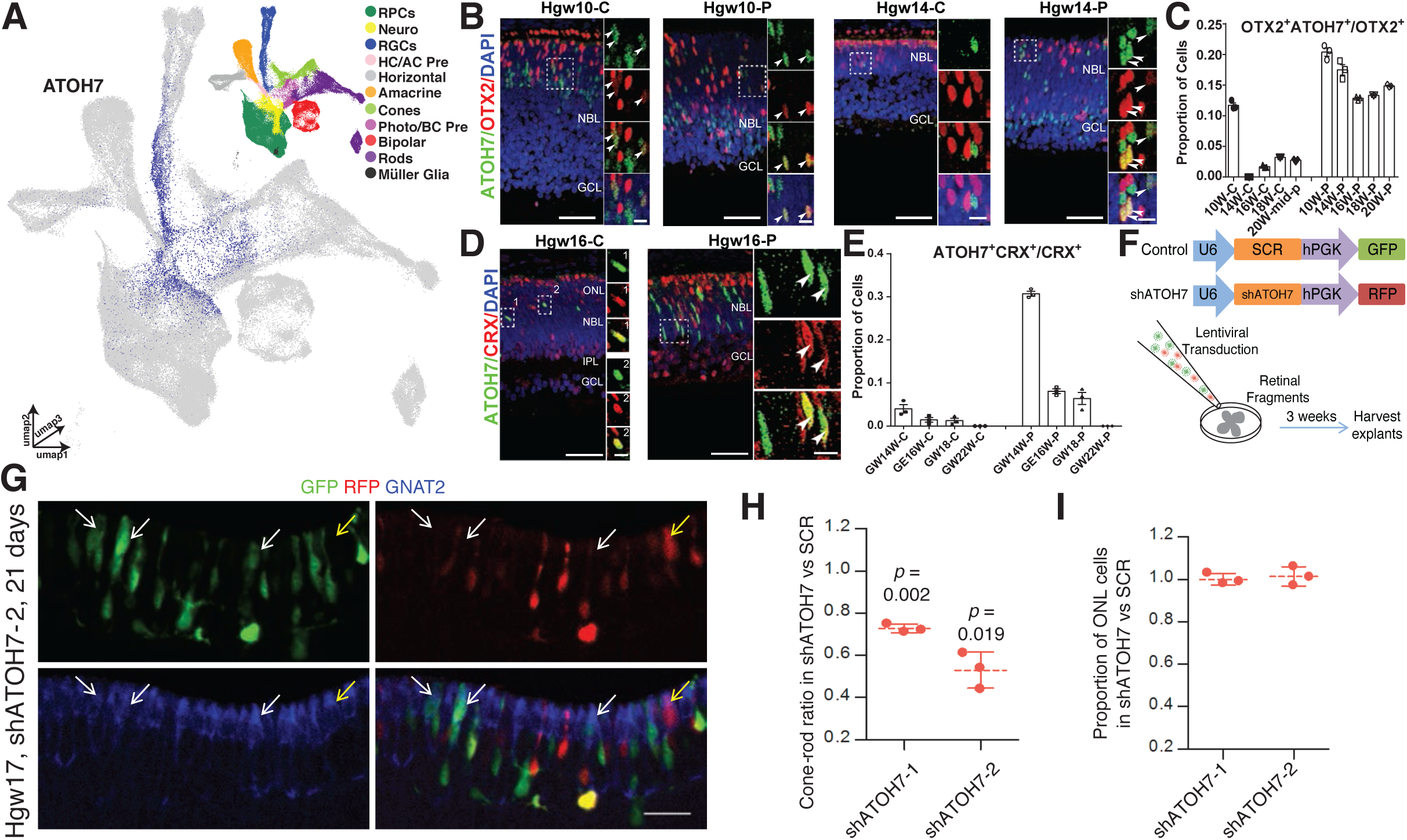
Knockdown of ATOH7 promotes differentiation of rod photoreceptor at the expense of late-born cones. (A) UMAP embedding of the retinal scRNA-Seq dataset, colored by relative expression of *ATOH7* and cell type annotation (top right). (B) Immunostaining for ATOH7 and OTX2 in central and peripheral Hgw10 and 14 retina, with zoomed in views of boxed regions. Scale bars: 50 μm and 10 μm (magnified views) (C) Quantification of OTX2+ cells that are also ATOH7*+* in central and peripheral retina at various ages. (D) Immunostaining for ATOH7 and CRX in the central and peripheral Hgw16 retina, with zoomed in views of boxed regions. Hgw16-C scale bar: 50μm (left), 5μm (right); Hgw16-P scale bar: 50μm(left), 10μm (right). Arrowheads indicate co-localization of markers and nuclei are counterstained with DAPI in panels B+D. (E) Quantification of CRX*+* cells that are also ATOH7*+* in central and peripheral retina at various ages. (F) Schematic diagram of *ATOH7* knockdown experiment. (G) Representative image from human retinal explants co-transduced with shSCR (GFP) and shATOH7-2 (RFP) lentiviruses and stained with cone marker GNAT2 (blue). White arrows, GNAT2+ cells expressing shSCR; yellow arrow, GNAT2+ cells expressing shATOH7. Scale bars, 20μm. (H) Ratio of cones/rods in shATOH7 vs shSCR cells. Data are presented as means ± SD. (I) Ratio of the ONL proportions in shATOH vs shSCR cells. Data are presented as means ± SD. Abbreviations: RPCs - retinal progenitor cells; Neuro - neurogenic cells; RGCs - retinal ganglion cells; HC/AC Pre - horizontal cell/amacrine cell precursors; Photo/BC Pre - photoreceptor/bipolar cell precursors; Hgw - human gestational weeks; GW - gestational weeks; C - central retina; P - peripheral retina; NBL - neuroblast layer; GCL - ganglion cell layer; ONL - outer nuclear layer; INL - inner nuclear layer; IPL - inner plexiform layer; C - central retina; P - peripheral retina.

These results imply that ATOH7 might act selectively in humans to promote the specification and/differentiation of cones in the peripheral retina. To directly test whether ATOH7 regulates human photoreceptor genesis, we co-transduced GW17-19 explants with lentiviral vectors that express control or ATOH7 shRNA together with GFP or RFP, respectively (Fig. 7F). Quantifying control and test shRNA transduced cells in the identical retinal fragment avoids the confounding effects of positional variation in developmental stage. We confirmed efficient depletion by shATOH7 vectors relative to a scrambled (shSCR) control (Fig. S7E,F). Explants were assessed 21-23 days after co-transduction. Cone and rod proportions were quantified in control and test cells using positive and negative labeling in the ONL for one of two cone markers (RXRγ or GNAT2) or a rod marker (NRL), and the cone-rod ratio in shATOH7 cells relative to shSCR cells was calculated. Irrespective of which of the three staining methods was used, depleting ATOH7 with either of two separate shRNAs reduced cones in favour of rods (Fig. 7G,H; Fig. S7G-J). Depleting ATOH7 did not affect overall photoreceptor numbers because the ratio of ONL proportions in shATOH7 vs. shSCR was 1 (Fig. 7I). Together, these data indicate that ATOH7 promotes cone genesis in the human fetal retina.

## Discussion

Our transcriptomic analysis using both embryonic stem cell-derived retinal organoids and primary tissue at single-cell resolution encompasses nearly the full time-course of human retinal development. While bulk RNA-Seq of the developing human retina and scRNA-Seq of human retinal organoids are already available, they have important limitations (Collin et al., 2019; Hoshino et al., 2017; Hu et al., 2019; Mellough et al., 2019)). Bulk RNA-Seq cannot resolve cell-type specific changes in gene expression. Furthermore, the extent to which retinal organoids fully recapitulate *in vivo* development at the individual cell level is still not entirely clear (Cowan et al., 2019). The combinatorial use of scRNA-Seq and human developing retinal tissue in our dataset overcomes some of these limitations. Combined, the dataset offers a valuable resource for identifying the gene regulatory networks that occur in human retinal development. Using this dataset in parallel with data previously generated from mouse (Clark et al., 2019; Lo Giudice et al., 2019) can highlight similarities and discrepancies across evolution, and help determine whether specific mouse models are actually relevant to human retinal disease. Furthermore, these findings open new avenues of research with respect to spatial patterning, cell fate specification, and the function of individual neuronal subtypes of the human retina.

This analysis identified some major differences between the gene expression pattern of immature cones analyzed in human retinal organoids and primary tissue (Fig. S3A-C). We hypothesize that this difference results from the abrupt changes in culturing conditions with the addition or subtraction of growth factors, signaling pathway inhibitors, or oxygen during the human retinal organoid culturing process. One of these treatments, in particular, is the addition of the gamma-secretase and Notch-pathway inhibitor DAPT from days 29-45 of organoid culture to assist with photoreceptor specification (Jadhav et al., 2006; Wahlin et al., 2017; Yaron, 2006). Interestingly, 92.7% (1591 of 1716) of organoid-derived cones in our dataset were derived from organoids within this treatment window (i.e. 30 or 42 days in culture organoids). This suggests that while the treatment of organoids with DAPT does stimulate photoreceptor specification, the differentiation trajectory of these organoid-derived cones varies markedly from that seen *in vivo*, even though more mature organoid-derived cones and native cones eventually show broadly similar gene expression profiles (Cowan et al., 2019).

Human retinal neurogenesis occurs over a much longer interval than in mouse. While many of the identified cell-type specific markers and differentially expressed genes across the developmental trajectory of each major retinal cell type are conserved in mouse, there are still some major differences. For example, *Clu* is specific to Müller glia in mice, but is expressed in both RPCs and Müller glia in human. Interestingly, a previous study has shown that *Clu*+ revival stem cells in the intestine are multipotent, capable of giving rise to the major cell types of the intestine and transiently expanding in a YAP1 dependent manner (Ayyaz et al., 2019). Since *Clu* is strongly expressed in both human RPC and Müller glia, it is possible that manipulating the YAP1 pathway will induce the regenerative pathway of human Müller glia. Consistent with this, two recent papers have implicated Hippo pathway signaling in regulating Müller glial quiescence, and YAP overexpression was sufficient to induce Müller glia to reenter the cell cycle (Hamon et al., 2019; Rueda et al., 2019). A number of genes, including several transcription factors such as *HES4* and *HKR*, show highly cell type-specific expression in human retina, but are absent from the mouse genome altogether. This raises the question of whether these genes are used in regulating human-specific aspects of retinal development, or have simply taken over evolutionarily-conserved functions carried out by related genes in mice.

The presence of the fovea is perhaps the most obvious anatomical difference between the retinas of primates and other mammals. We found differentially expressed genes between peripheral and macular RPCs and Müller glia, allowing us to elucidate the molecular underpinnings of cone-rich foveal formation. One of the genes examined, *CYP26A1* (retinoic acid-metabolizing enzyme), is conserved in chicken, and is important for creating a rod free zone by catabolizing retinoic acid (da Silva and Cepko, 2017). This suggests that conserved mechanisms inhibiting the retinoic acid signaling pathway are involved in the formation of a high-acuity, cone-rich, rod-free zone. Other genes that are strong candidates for controlling foveal development that are both enriched in these cells and not strongly positively correlated with pseudotime include *DIO2*, which promotes differentiation of L/M cones at the expense of S-cones (Eldred et al., 2018), the Wnt pathway inhibitors *SFRP2* and *FRZB*, and secreted antiproliferative factors such as *CTGF* and *PTN.* Wnt signaling acts during early stages of retinal development to inhibit central retinal identity (Cho and Cepko, 2006; Liu et al., 2007), and its presence here suggests that it may play an analogous role in control of foveagenesis. Antiproliferative factors such as *CTGF* and *PTN* may mediate the early cessation of proliferation seen in the foveal region (Diaz-Araya and Provis, 1992; van Driel et al., 1990), although the fact that *CTGF* expression is also enriched foveal Muller glia in adult macaque suggests that it may also have other functions related to controlling mature glial function or morphology (Peng et al., 2019). The coordinated action of these signaling pathways in RPCs may provide the initial signals that pattern the fovea. Finally, macular-enriched expression of a number of genes encoding cell adhesion and extracellular matrix components *(ITGB1, COL1A2, ITM2B*) is observed in RPCs and MG, suggesting that these may be candidates for morphogenesis of the foveal pit. Although macular-enriched genes that are not positively correlated with pseudotime are found that are enriched in multiple neuronal cell types, including several that are enriched in both developing photoreceptors of the human macula and adult macaque fovea (Peng et al., 2019), their function is less clear, and awaits direct functional analysis.

Similar to previous studies in macaque and adult humans, we also observed two subtypes of horizontal cells in human retina, consistent with those classified in previous studies of macaque retina (Peng et al., 2019). Although morphological studies of adult human retina have identified three distinct subtypes of horizontal cells (Kolb et al., 1994), we only observed clear evidence for two. In humans, these two subtypes express different combinations of neurotransmitter receptor genes. Most prominently, *LHX1*+ horizontal cells specifically express dopaminergic receptor *DRD2* while no other dopaminergic receptor genes are expressed in either horizontal cell subtypes. While dopaminergic amacrine cells are thought to be the only retinal cells that produce dopamine, we detected weak expression of the dopamine transporter *SLC6A3* in both rods and bipolar cells. The expression of *DRD2* on *LHX1+* horizontal cells and potential dopamine transporter transcript expression in rods and bipolar cells, therefore, implies that *LHX1+* horizontal cells may selectively respond to a dopamine signal within the rod spherule.

We also observed that *ISL1+/CALB1+*, unlike *Lhx1+*, horizontal cells lack *MEGF11* expression. MEGF11 regulates homotypic repulsion of horizontal cells in mouse (Kay et al., 2012). Since retinal subtypes are frequently spaced evenly across the retina, but randomly positioned with respect to other subtypes (Rockhill et al., 2000), it will be interesting to identify the determinants of *ISL1+* horizontal cell retinal mosaic spacing independent of *MEGF11* and define these novel mechanisms of horizontal cell homotypic repulsion in the developing human retina. Other evolutionary differences we observe include the re-emergence of *LHX1+* and *CALB1+* horizontal cells in primate retinas (Peng et al., 2019), whereas the genes are co-localized in the single type of mouse horizontal cell. Since chickens also possess distinct populations of *Isl1+* and *Lhx1+* horizontal cells, with two morphologically distinct populations of *Isl1*+ cells (Suga et al., 2009), we hypothesize that nocturnal rodents consolidated these populations to one cell type. Further examination of number of cellular subtypes within the retina across evolutionarily divergent species will address whether or not the number of horizontal cell subtypes directly correlates with the number of cone subtypes.

To examine gene regulatory networks in an unbiased manner and without *a priori* knowledge of gene interactions, we used scCoGAPS pattern identification and ProjectR latent transfer learning. These tools provide a robust, transferable way for comparing gene-coexpression networks across species (Sherman et al., 2019; Stein-O’Brien et al., 2019). These techniques identified patterns of gene expression that were both shared and divergent across species. Furthermore, these analyses highlighted genes such as *LOXL1*, implicated in pseudoexfoliation syndrome (Thorleifsson et al., 2007), to have distinct expression patterns across human and mouse retinal development. This not only implies that pseudoexfoliation syndrome may present with a retinal pathology during development, but also highlights some of the unexpected pitfalls of modelling human diseases in mice. Divergent patterns between the two species also raises the question of what are the evolutionary drivers behind these divergences. In human corticogenesis, it has been hypothesized that coordinated changes in regulatory domains of co-expressed genes drive cortical evolution (Reilly et al., 2015). This is further highlighted by large transcriptional differences observed in brain cell type evolution across the primate lineage, despite little sequence divergence in protein coding genes (Khrameeva et al., 2019). Hence, it is possible that the divergent expression patterns of conserved genes across human and mouse retinal development corresponds to changes in the use and/or sequence of transcriptional regulatory elements.

Finally, we examined the role of *ATOH7* within late developmental periods of retinogenesis. Patients with mutations within the *ATOH7* coding or regulatory sequences exhibit persistent fetal vasculature (PFV) and/or nonsyndromic congenital retinal nonattachment (NCRNA), respectively (Ghiasvand et al., 2011; Prasov et al., 2012). Both of the diseases, however, are associated with defects in RGC development. Here, we show *ATOH7* is also involved in the specification of late-born cones, suggesting alterations in the proportion of cones to rods may also be observed in patients with either PHPV or NCRNA. *Atoh7* mutant mice generate additional cones at the expense of RGCs (Brown et al., 2001), although *atoh7* zebrafish mutants retinas exhibit no change in cone development (Kay et al., 2001). Together, these data suggest *ATOH7* may have different mechanisms of function in retinal cell fate determination across evolution. The exquisite sensitivity or resistance of RB1-deficient human or murine cones to tumorigenesis reflects the intrinsically high expression of oncogenes such as MDM2 and MYCN specifically in human cells (Xu et al., 2009). Our results highlight a fundamental difference between the mechanisms by which human and mouse cones are specified, which presumably influences the resulting cone circuitry and, therefore, susceptibility to *RB1* loss.

The rapid improvement of techniques for the culturing of human ES and iPS-derived retinal organoid preparations makes it feasible to directly study the functional role of candidate extrinsic and intrinsic regulators of human retinal development identified in this study. While functional studies using human retinal organoids have thus far largely focused on analysis of evolutionarily conserved gene regulatory networks (Eldred et al., 2018), they provide an accessible and tractable system for studying aspects of retinal development that cannot be readily modelled in mice and early fetal development. The combinatorial application in organoid preparations of growth and differentiation factors identified in our study has the potential to highlight molecular signals that are sufficient to induce foveagenesis. Coupled with the use of CRISPR-mediated knockout approaches, this will lead to direct functional tests of human retinal genes that lack mouse orthologs or show markedly different expression patterns in developing retina.

## Acknowledgements

We thank J. Nathans, A. Kolodkin, R. Johnston Jr., S. Chen, and P. Ruzycki for comments on the manuscript. We thank the Transcriptomics and Deep Sequencing Core at Hopkins for assistance in sequencing and Dr. Jeff Wrana and Daniel Trcka for technical assistance with preparing single cell libraries and both the Morgentaler Clinic in Toronto and the donors for access to developing retinal tissue.

## Material and Methods

### STAR⋆Methods

#### Key Resources Table

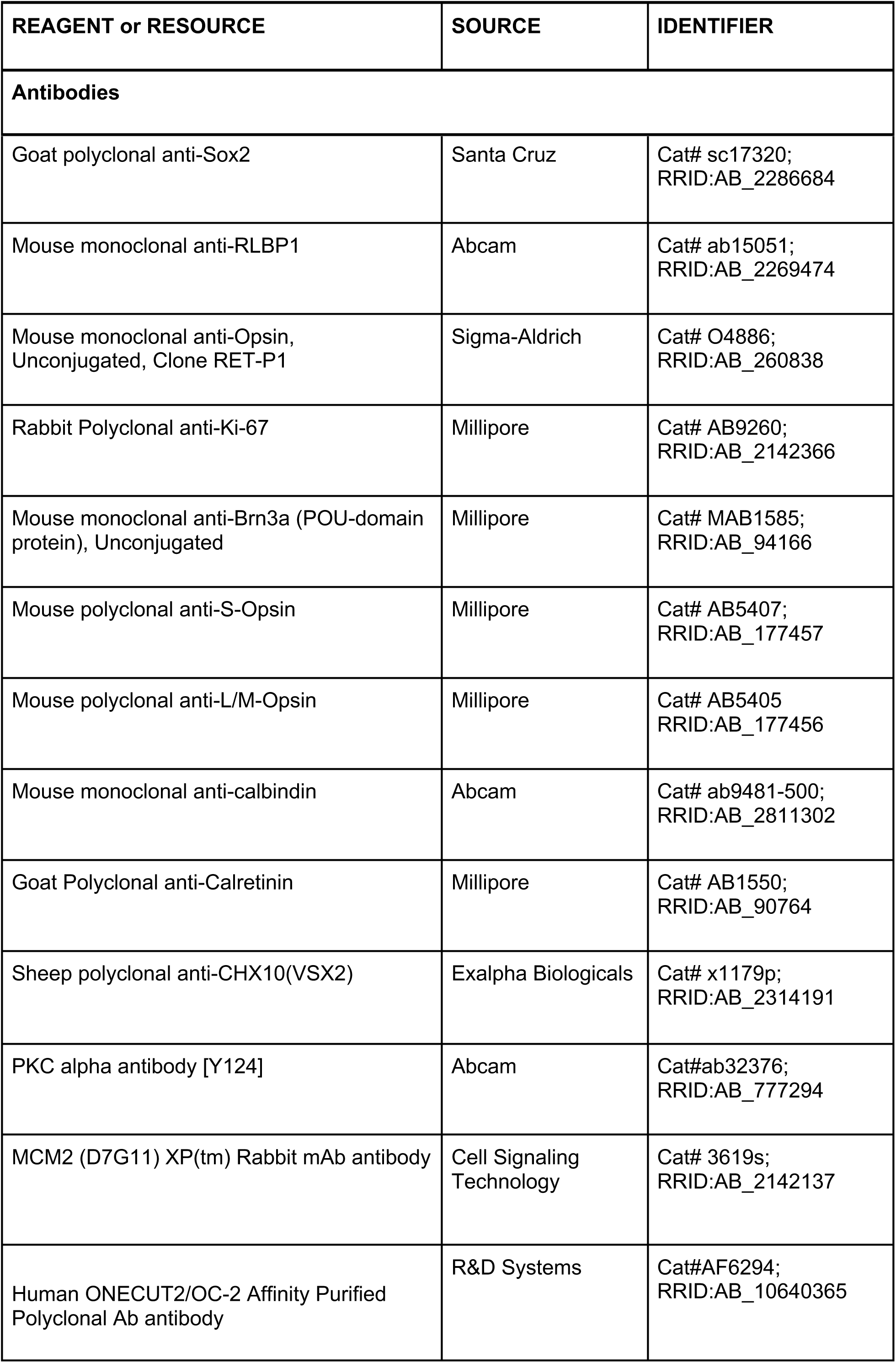
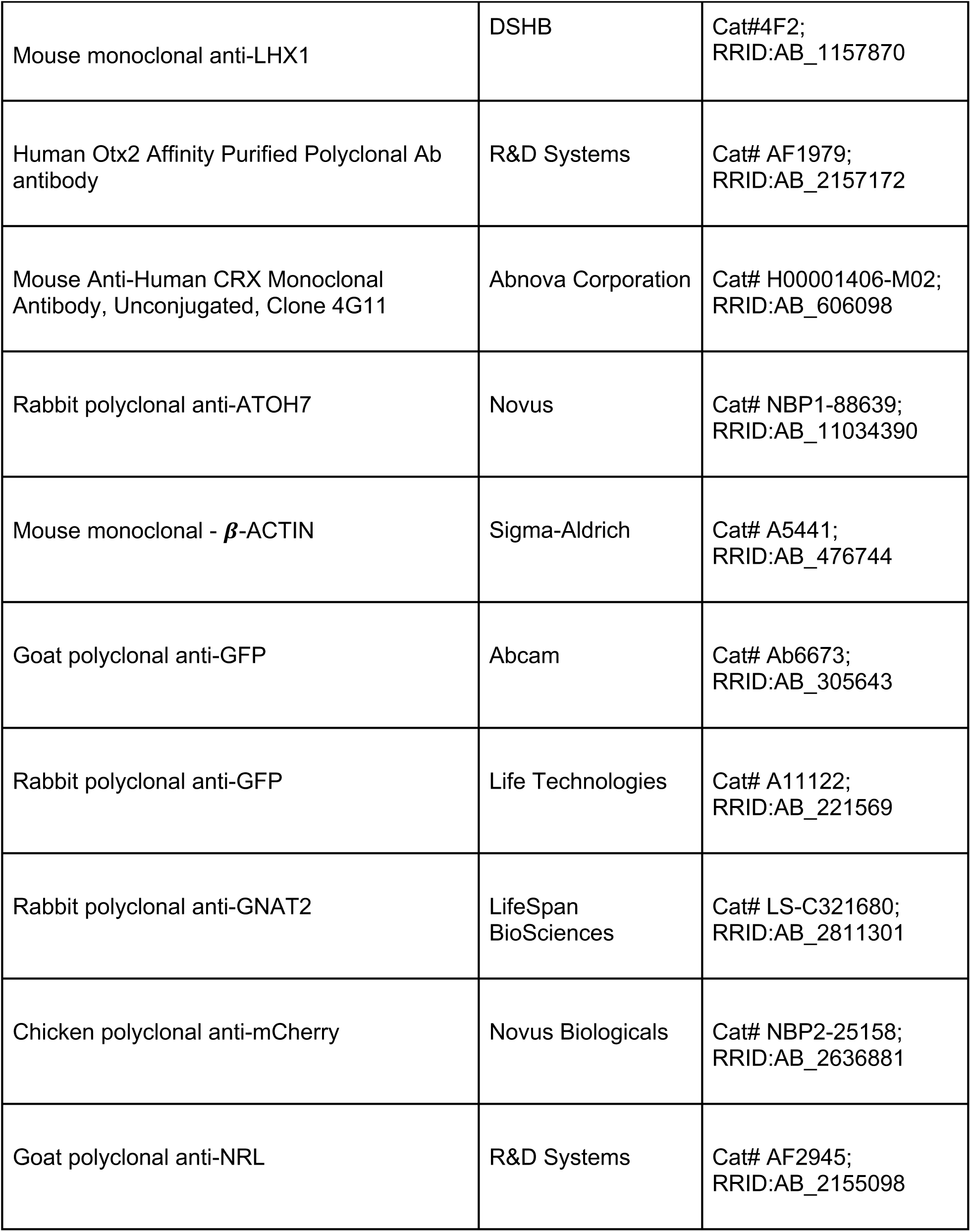
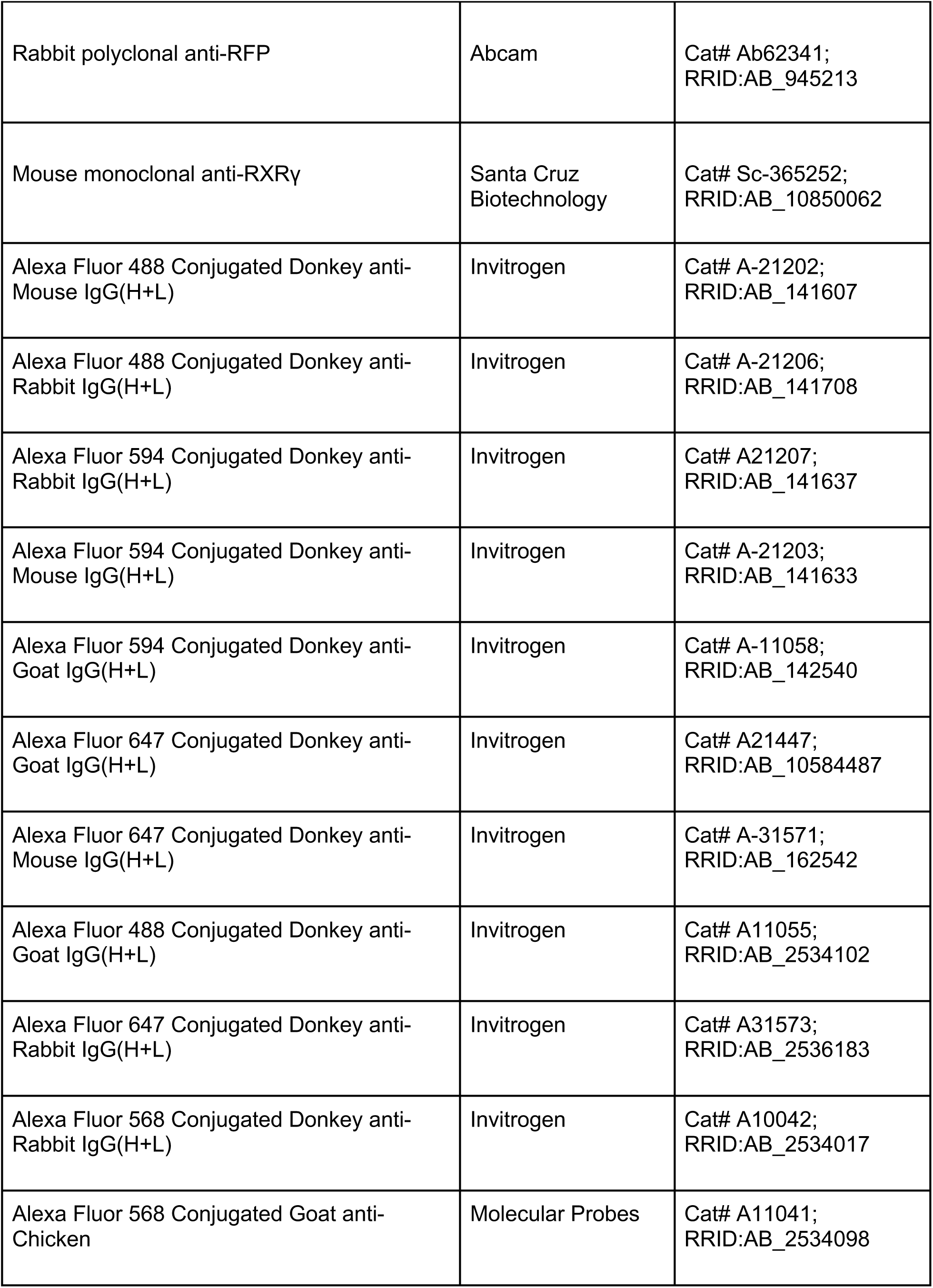
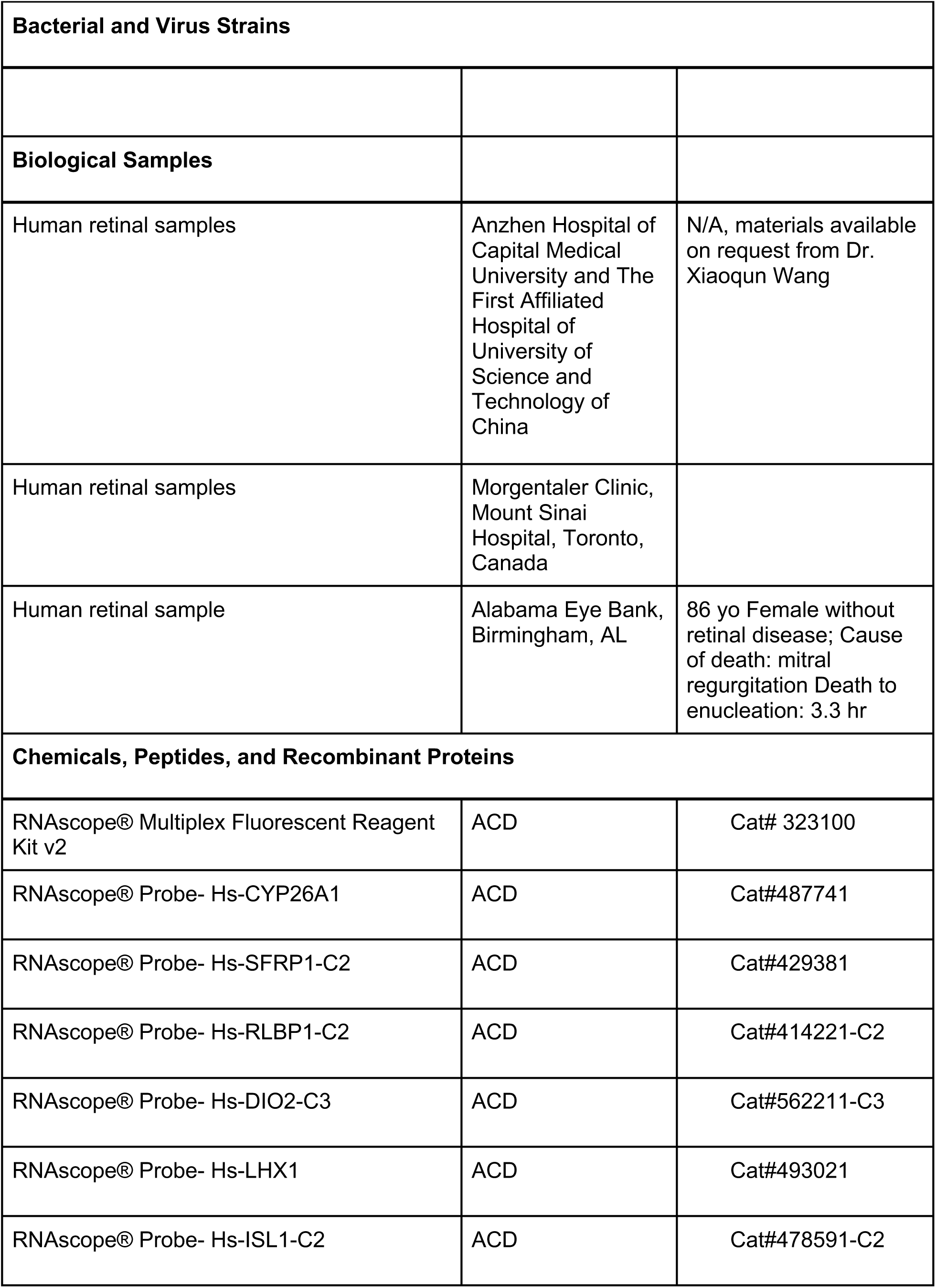
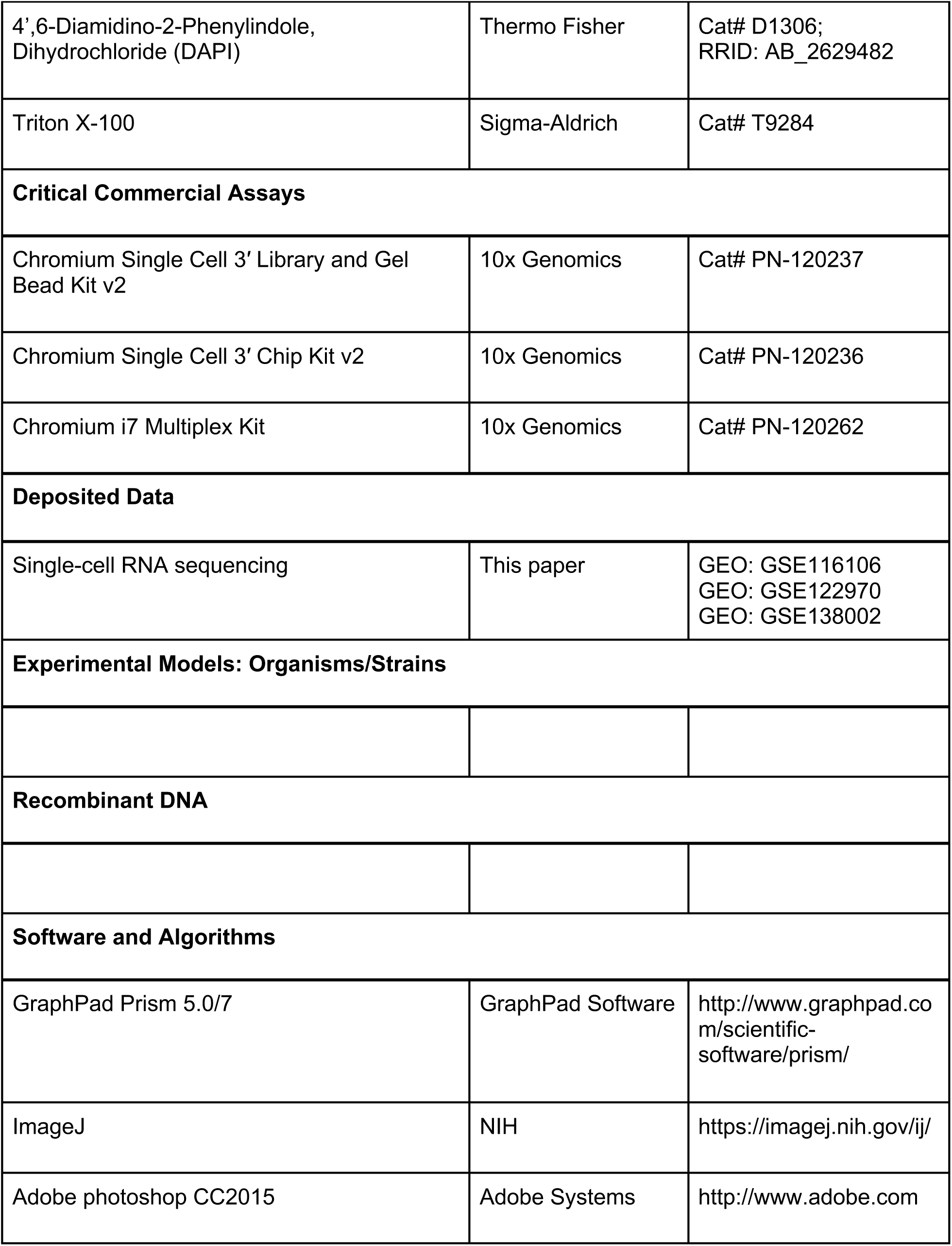
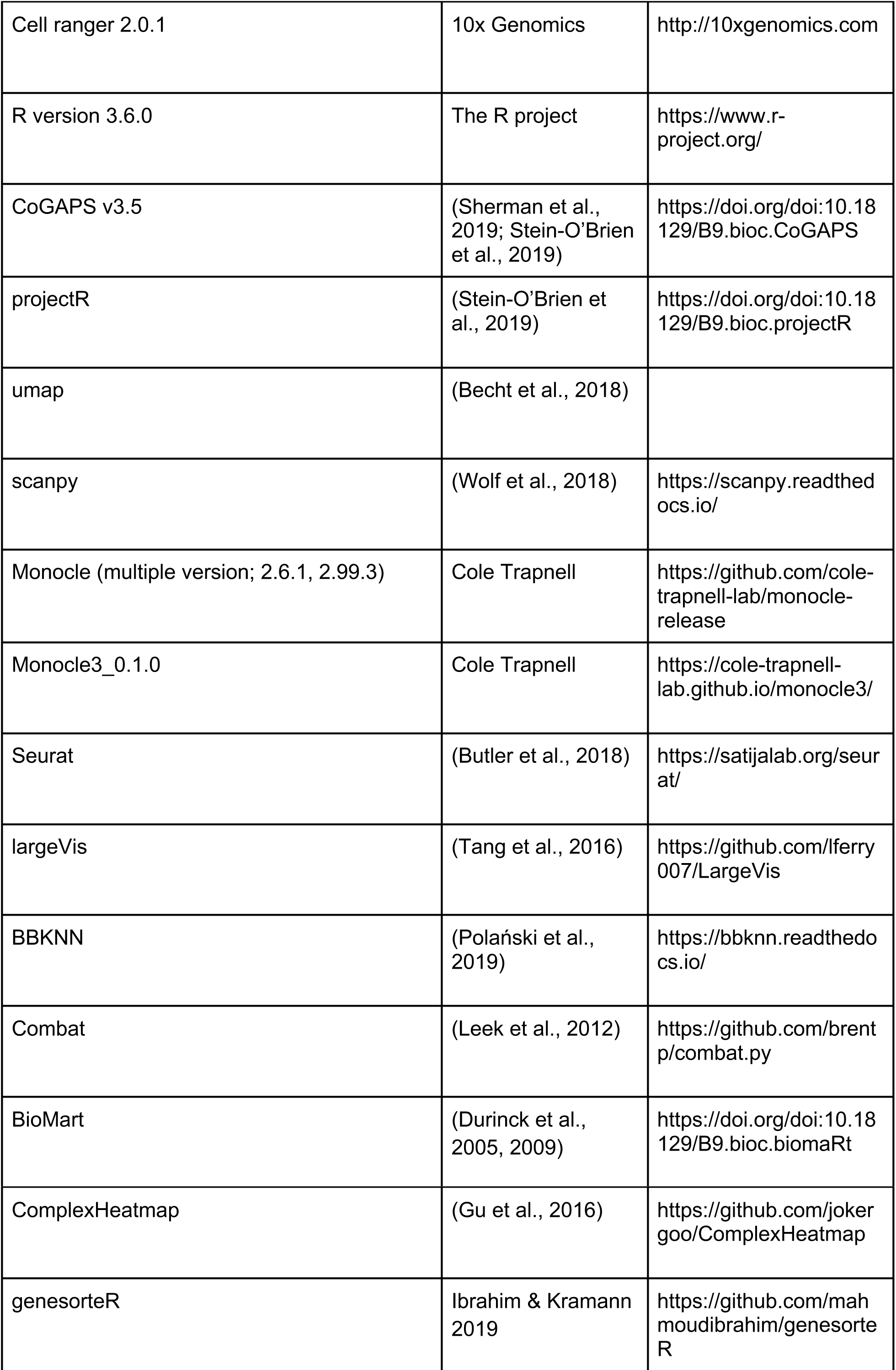

#### Ethics Statements

##### Canadian Samples

Retinas were obtained from the Morgentaler Clinic in Toronto with approval from the Research Ethics Board (REB, REB13-0132-E) of Mount Sinai Hospital in Toronto, Canada. All donors read the consent form approved by the REB before surgical procedures, and voluntarily donated developing eye samples. The gestational age was estimated by a combination of clinic intakes, ultrasound, crown-rump, and foot length measurements where possible (FitzSimmons et al., 1994; Shepard, 1975). Eye samples collected were held on ice for up to 6 hour in the retina culture medium, which is IMDM with 10% FBS and 1X Antibiotic-Antimycotic (Life Technologies, Cat#15240062).

##### Chinese Samples

The human tissue collection and research analysis was approved by the Reproductive Study Ethics Committee of Beijing Anzhen Hospital and The First Affiliated Hospital of University of Science and Technology of China. The informed consent forms were designed under ISSCR guidelines for human developing tissue donation and were in strict observance of the legal and institutional ethical regulations for elective pregnancy termination specimens at Beijing Anzhen Hospital and The First Affiliated Hospital of University of Science and Technology of China. All protocols were in compliance with the ‘Interim Measures for the Administration of Human Genetic Resources’ administered by the Ministry of Science and Technology of China.

##### US Sample

The use of human globes obtained from Tissue Banks was approved by the Johns Hopkins Institutional Review Board.

#### Preparation of Retinal Organoids

Retinal organoids were prepared as described (Wahlin et al., 2017) from induced pluripotent stem cells (iPSCs) derived from the IMR-90 (ATCC) cell line. On Day 0, iPSCs were plated at 3,000 cells/well in 96-well U-bottom plates for forced aggregation into embryoid bodies, cultured in mTeSR1 medium (Stem Cell technologies) + 5μM blebbistatin in hypoxic conditions (5%O_2_/10%CO_2_). Cells are then transferred to normoxia (20%O_2_/5%CO_2_) after 24 hours and medium is gradually (Days 1-7) changed to BE6.2 (10ml E6 stock (97mg insulin 53.5mg holo-transferrin, 230mg L-ascorbic acid, 5μl 14mg.ml sodium selenite, to 100ml with H_2_O - of note, no NaHCO_3_ was used in E6 stocks), 5.0ml B27 (without vitamin A), 2.5ml Glutamax (100X), 2.5ml NEAA (100X), 2.5ml Pyruvate (100x) 1.0ml NaCL (21.9g/.1L) and DMEM to 250ml) + 1%(v/v) matrigel supplemented with 3uM Wnt inhibitor, IWR-1. Medium exchange on Days 8 and 9 in culture removes the WNT inhibitor IWR-1 and matrigel. On Day 10, organoids are washed with HBSS and transferred to 10cm dishes and feed with BE6.2 medium + 100nM Smoothened agonist SAG. Optic vesicles are manually excised from organoids between Days 10 and 14. From Days 12-18 medium is changed every other day with LTR (125ml F12, 50ml FBS, 10ml B27, 5ml Glutamax (100X), 5ml NEAA (100X), 5ml pyruvate (100X), 500 μl taurine (1000X - 1M stock), DMEM to 500ml) medium + 100nM SAG. On Day 18, culture medium is changed to LTR medium without SAG. To promote retinal differentiation and maturation, medium is changed to LTR medium supplemented with 500nM all-*trans* retinoic acid (ATRA). From Days 28-42, LTR medium is supplemented with both 500nM ATRA and 10μM DAPT.

#### Retinal Dissociation for single cell sequencing

##### Canadian Samples

Developing eyes at gestational ages week 9 to week 19 were sterilized in 70% ethanol for 3 seconds, rinsed twice in cold phosphate-buffered saline (PBS) (Wisent Bioproducts, Cat#311-010-CL), and transferred to IMDM for retinal dissection. Retinas were dissociated with Papain Dissociation System (Worthington Biochemical, Cat# LK003150). Briefly, retinas were incubated in papain solution for about 15 minutes at 37°C and 5% CO2, with gentle pipetting every 5 minutes. With dissociation to ∼20-cell clusters, cells were suspended in 10 volumes of sterile phosphate buffered saline (PBS), pelleted by centrifugation at 300 x g for 10 minutes, and re-washed with PBS, followed by digestion with 0.05% trypsin/EDTA (Wisent Bioproducts, Cat# 325-042-EL) with gentle pipetting to produce single cell suspension, washed with 10 volumes of retina culture medium, and resuspended in proper volumes of the culture medium for single cell sequencing procedure. Cell viability (>90%) was confirmed by negative staining with trypan blue (Life Technologies, Cat#15250061). Single cell libraries were then prepared using the 10x Single Cell 3’ v2 Reagent Kits according to the manufacturer’s instructions and sequenced on an Illumina NextSeq500 using recommended sequencing parameters (Read 1 - 26bp; Read 2 - 98bp; i7 Index - 8bp; i5 Index - 0).

##### Chinese Samples

Gestational age was measured in weeks from the first day of the woman’s last menstrual cycle to the sample collecting date. Developing retinal tissue samples were collected in ice-cold artificial cerebrospinal fluid (ACSF) containing 125.0 mM NaCl, 26.0 mM NaHCO_3_, 2.5 mM KCl, 2.0 mM CaCl_2_, 1.0 mM MgCl_2_, 1.25 mM NaH_2_PO_4_; pH 7.4, bubbled with 95% O_2_/5% CO_2_. The retina macula samples (2mm) were defined by lack of vascular. Whole retinal samples were gently separated into small pieces and then centrifuged at 200g for 2min. The supernatant was removed and 500ul digestion buffer (2mg/ml collagenase IV (Gibco), 10 U/μl DNase I (NEB), and 1mg/ml papain (Sigma) in PBS) was added. The tissue was then rotated and incubated at 37°C on a thermo cycler with 300g for 15-20 min. Samples were triturated every 5 min to digest the tissue sample into single cell suspensions. Finally, inactivation of enzymatic digestion was induced through the addition of an equivalent volume of 10% fetal bovine serum (Gibco) in PBS. Cells were pelleted and resuspended in 0.04% BSA/PBS and stained with 7-amino-actinomycin D (7-AAD) for 10 min on ice in order to check for cell viability. 7-AAD-negative cells were collected by FACS and resuspended in 0.04% BSA/PBS. Single cell libraries were then prepared using the 10x Single Cell 3’ v2 Reagent Kit according to the manufacturer’s instructions and sequenced on an Illumina HiSeq4000 with 150bp paired-end reads.

##### US Samples

Retinal organoids were processed as described in Clark *et al*., 2019. Multiple organoids were pooled and placed in 200μl of cold HBSS per organoid, with an equivalent amount of Papain dissociation solution (for 1ml - 100μl freshly prepared 50mM L-Cysteine (Sigma), 100μl 10mM EDTA, 10μl 60mM 2-mercaptoethanol (Sigma), and Papain added to 1mg/ml (Worthington), to 1ml with reagent-grade water). Organoids in dissociation solution were then placed at 37C for 10 minutes, with slight trituration every 1-2 minutes. Enzymatic digestion was halted with addition of 600μl Neurobasal Medium supplemented with 10% FBS for every 400μl of HBSS/dissociation solution. 5μl/organoid DNaseI (RNase free Recombinant DNaseI; Roche) was added and incubated 5-10 minutes at 37C. The dissociation solution was then gently triturated using a P1000 pipette tip and cells were pelleted through centrifugation for 5 minutes at 300RCF. Supernatant was carefully aspirated off the cell pellet, followed by resuspension of the cellular pellet in 1-5ml Neurobasal media with 1% FBS. Cellular aggregates were removed by straining cells through a 50μm filter. Single-cell libraries were then prepared using the 10x Single Cell 3’ v2 Reagent Kits according to the manufacturer’s instructions and sequenced on an Illumina NextSeq500 using recommended sequencing parameters (Read 1 - 26bp; Read 2 - 98bp; i7 Index - 8bp; i5 Index - 0).

##### Human adult retinal sample

A human globe from an 86 year old Caucasian female who died of a myocardial infarction and had no known ocular disease other than cataracts, was obtained from the Alabama Eye Bank (Birmingham, AL) and processed within 3.3 hours after death. The study was approved by the Johns Hopkins Institutional Review Board. To dissect the neural retina. the anterior segment was first removed by incising the scleral behind the limbus, to remove the anterior parts, lens and vitreous body. The neural retina was then peeled off from the eyecup, and retinal cells were dissociated using Papain Dissociation System (Worthington Biochemical, Lakewood, NJ) following the manufacturer’s instructions. Dissociated cells were resuspended in ice-cold PBS, 0.04% BSA and 0.5 U/UL of RNAse inhibitors. Cells were then filtered through a 50um filter.

#### Library preprocessing

Resulting sequencing outputs were processed through the CellRanger2 mkfastq and count pipelines using default parameters. Transcript reads were quantified using the 10x Genomics Human reference index (refdata-cellranger-GRCh38-3.0.0). Cells were then given unique, sample-specific cell identifiers to prevent duplicate cell names in the aggregate dataset from re-use of barcodes across samples.

#### Aggregation of Datasets, initial processing and cell type assignment

##### Human retinal organoids

Resulting count matrix files from Cellranger alignments/counting of retinal organoids were imported and aggregated in Monocle2 R/Bioconductor (Qiu et al., 2017). First, cells with > 40,000 Total_mRNAs were removed as outliers. We then identified genes with high biological coefficient of variation by first normalizing sequencing depth across all organoid-derived cells using the Waddington-OT transformation to transcript copies per 10,000 (CPT) (Schiebinger et al., 2019) and then using a generalized additive model (MGCV R package; (Wood et al., 2015)) fit to the log2 mean CPT versus a cubic spline fit to the log2 coefficient of variation across all genes with detectable expression in >10 cells. Transcripts that displayed >1.1 residual to the fit were chosen as ‘high-variance’ genes. Dimension reduction was then performed on the resulting 2441 high variance genes (Table S1) using the first 24 principal components as input into largeVis dimension reduction (Tang et al., 2016). Cell type annotation of Retina/RPE versus non-Retina was performed using eye-field marker genes including *RAX, PAX6,* and *VSX2* while excluding markers of ventral telencephalon and hypothalamus, including *NKX2.1*, *DLX5* and *DLX6*. The resulting 11,758 cells Retinal/RPE cells of the original 25,461 organoid cells were then used for aggregation with the *in vivo* samples.

##### Human retinas

All 20 samples of primary retinal tissue were manually aggregated from the individual Cellranger matrix files, resulting in 113,999 individual cells. Initial processing proceeded as above with the following differences: 1) Cells with >10,000 Total_mRNAs were removed, 2) A residual cutoff for determination of high variance genes was set at 1.15, resulting in 2719 highly variable genes that were used for input into dimension reduction, 3) the first 17 dimensions were used as input to PCA, and 4) dimension reduction was performed using UMAP within the ‘reduceDimension’ function of Monocle3alpha, with the following parameters: max_components = 3, metric= ‘canberra’, min_dist = 0.34, n_neighbors = 50, residualModelFormulaStr = “∼Total_mRNAs + sample”, random_state = 123456L. The resulting 3-dimensional structure was clustered using the ‘clusterCells’ function of Monocle3alpha with the following parameters: use_pca = FALSE, k=15,res=4.0e-4, method = ‘louvain’, gaussian = TRUE, louvain_inter=5, set.seed(123456). Preliminary cell type annotation was performed based on cluster markers from the ‘find_cluster_markers’ function and examination of expression of known marker genes within the individual clusters. Non-retinal derived cells (microglia, vasculature, *etc.*) and annotated doublets (identified through incompatible gene expression) were removed from the cell dataset at this time, resulting in an accumulated cell total of 107,013 *in vivo* retinal cells.

##### Final aggregation

Cell datasets from the organoids and *in vivo* retinas were merged to create a single cell dataset. We again removed cells with >10,000 Total_mRNAs, thus reducing the final dataset to 11,542 organoid-derived cells and 107,013 human retina-derived cells (118,555 total cells). The same exact dimension reduction parameters were used for the total aggregate as the *in vivo* retinas, using the high variance genes from the *in vivo* developing retina dataset. Clustering on the 3-dimensional UMAP was performed using ‘clusterCells’ function of Monocle3alpha using the following parameters: use_pca = FALSE, k=15,res=1.0e-3, method = ‘louvain’, gaussian = TRUE, louvain_inter=5, set.seed(123456). Determination of corresponding cell type of clusters was performed based on marker gene expression as previously described. Cell cycle phase of primary and neurogenic RPCs was determined using the ‘CellCycleScoring’ function of Seurat.

#### Immunohistochemistry of retinal tissue

Human retina tissue samples were fixed overnight in 4% paraformaldehyde. The fixed retinas were dehydrated in 20% and 30% sucrose in PBS at 4°C and embedded in optimal cutting temperature medium (Thermo Scientific). Thin 20-40μm cryosections were collected on superfrost slides (VWR) using a Leica CM3050S cryostat. For immunohistochemistry, antibodies against the following proteins were used at the following dilutions: Goat anti-SOX2 (1:250, Santa Cruz), Rabbit anti-Mki67 (1:200, Milipore), Mouse Anti-Brn3a (1:500, Millipore), Mouse anti-RLBP1 (1:500, Abcam), Mouse anti-Rod-OPSIN (1:1000, Sigma), Rabbit anti-S-OPSIN (1:500, Millipore), Rabbit anti-L/M-OPSIN (1:500, Millipore), Mouse anti-Calbindin (1:500, Abcam), Rabbit anti RRKCA (1:500,Abcam), Sheep anti VSX2 (1:400, Exalpha Biologicals), Rabbit anti MCM2(1:200 Cell Signaling), Sheep anti ONECUT2 (1:500 R and D Systems), Mouse anti LHX1(1:200 DSHB), Rabbit anti ISL1(1:200 Abcam), Goat anti OTX2 (1:200 R and D Systems), Mouse anti CRX(1:200 Abnova Corporation), Rabbit anti ATOH7(1:200 Novus) and Goat anti-Calretinin (1:500, Millipore). Primary antibodies were diluted in blocking buffer containing 10% donkey serum, 0.2% Triton X-100 and 0.2% gelatin in PBS at pH 7.4. Alexa Fluor 488, Alexa Fluor 594 or Alexa Fluor 647 fluorophore-conjugated secondary antibodies (1:500) (Life Technologies) were used as appropriate. Cell nuclei were stained with DAPI (1:10000). Images were collected using an Olympus FV1000 confocal microscope.

#### RNAscope

RNAscope® detection was performed in strict accordance with the ACD RNAscope® protocol (Wang et al., 2012). Briefly, retina sections were dehydrated in sequential incubations with ethanol, followed by 30 min Protease III treatment and washing in ddH2O. Appropriate combinations of hybridization probes were incubated for 2 hours at 40°C, followed by fluorescence labeling, DAPI counterstaining, and mounted with Prolong Gold mounting medium.

#### Diffusion Pseudotime & Trajectory Analysis

We exported the cell dataset matrices used in Monocle to create an object to perform pseudotime in Scanpy v1.4 (Wolf et al., 2018).The dataset was subset to relevant cell types based on the desired analysis, with cells from samples displaying significant batch effects (Hgw12) or discontiguous trajectories in UMAP dimension reductions – likely resulting from gaps in sampling ages (Hpnd8 and Adult) – removed from the analyses. High variance genes used as input to Scanpy are listed in Table S2 for each cell type analysis. Diffusion Pseudotime was used to reconstruct the trajectory from least mature cells to final cell fate(s) as previously described (Haghverdi et al., 2016; Wolf et al., 2018). Batch effect corrections, if necessary, were conducted prior to determination of highly variable genes, unless the BBKNN test was used (Polański et al., 2019). The parameters used for assigning pseudotime values within Scanpy are listed in Table S3.

Corresponding pseudotime values from scanpy were assigned to cells within the Monocle cell dataset. Differential genes across pseudotime were assessed using the ‘differentialGeneTest’ function in Monocle, requiring that all differential transcripts being expressed in >= 10 cells. The following parameters were used for the differential gene test:

1. if one terminal cell fate, fullModelFormulaStr= “∼sm.ns(Pseudotime,df=3)”
2. if multiple terminal cell fates, fullModelFormulaStr = “∼sm.ns(Pseudotime,df=3)*Branch” and reducedModelFormulaStr = “∼sm.ns(Pseudotime,df=3)”.

If needed, known batch effects were included in the differential gene test parameters as previously described (Qiu et al., 2017; Trapnell et al., 2014).

For analyzing the trajectories from retinal progenitor cells to Müller Glial or Neurogenic Cells, cell cycle was regressed from the expression matrix using Seurat v3.0 to assign Cell Cycle Scores and Phase as previously described (Butler et al., 2018). These scores and phase were assigned to corresponding cells within the Monocle cell dataset. Cell Cycle Scores were regressed within the ‘preprocessCDS’ function (Qiu et al., 2017; Trapnell et al., 2014).

Differential gene expression across pseudotime analysis was then conducted on the regressed expression matrix as described above.

#### Annotation of retinal progenitor cell subtypes

Examination of the density of RPCs across pseudotime identified two clear density peaks. We annotated early versus late RPCs based on positioning across the pseudotime axis in relation to the trough between the peaks (dotted line - Fig. S2A; pseudotime value of 0.4609), with early RPCs corresponding to pseudotime values ≤ 0.4609 and late RPCs corresponding to pseudotime values >.04609.

#### scCoGAPS pattern discovery

##### Human Patterns

scCoGAPs analysis was performed on all retinal cells excluding the adult retinal samples. The large expression matrix size (33,694 genes by 118,555 cells) was subset down to a set of 3113 (Table S2) highly variable genes that excluded both mitochondrial and ribosomal protein coding genes. This down-sampling of genes allowed both faster implementation of scCoGAPS and was implemented to reduce the number of patterns that highlight sample batch effects of cellular stress or read depth. The log2 transformation of the expression matrix was used as input into the scCoGAPS function from CoGAPS v3.5 as previously described (Sherman et al., 2019; Stein-O’Brien et al., 2019). The parameters were default singleCell parameters, except nPatterns=100, nIterations=500, sparseOptimization = True, seed =830, nSets=27 (∼3960 cells/set) with annotation weights set to sample underrepresented cell types more.

##### Mouse Patterns

We used scCoGAPS from CoGAPS v3.5 to reanalyze the mouse retinal development dataset from Clark *et al.,* 2019, but limited the dataset to include only the retinal cells. We used 3164 high variance genes across the dataset, again excluding mitochondrial and ribosomal genes and used nSets = 25 (4414 cells/set).

#### Comparisons of pattern usage across species

To examine the extent to which human or mouse patterns corresponded to similar biological processes in the opposite species, we used projectR to project the log2 of the scRNA-Seq expression matrix into the scCoGAPS pattern amplitude matrix (Stein-O’Brien et al., 2019). The Ensembl database from BioMart was used to identify the corresponding homologs of the pattern genes (Durinck et al., 2005, 2009). As some mouse genes had multiple human homologs, the pattern weights of each of these human orthologs are combined by taking the maximum value. If a human gene had multiple mouse orthologs, the same gene weights were reused between the mouse orthologs.

The same process was used to determine mouse patterns and the projections of their use in human data matrix, except 3164 highly variable mouse genes were used, nSets=25 (4414 cells/set), and gene names were capitalized to find human orthologs.

#### Construction of shATOH7 lentiviral vectors

To suppress ATOH7 expression in developing human retina, pLKO.1-TRC cloning vector (Moffat et al., 2006) was purchased from Addgene (Cat#10878). The puromycin selection marker in the vector was replaced with GFP-P2A-Puromycin or RFP-P2A-Puromycin ORF cassette with Gibson Assembly Master Mix (New England Biolabs, Cat#E2611L) to generate pLKO.1-GFP-P2A-Puro or pLKO.1-RFP-P2A-Puro cloning vector. Specific cloning details are available on request. The target sequence for scrambled shRNA is CCTAAGGTTAAGTCGCCCTCG available from Addgene (Cat#1864) (Sarbassov et al., 2005), and the sequences of the hairpin pair are as follows:SCR-F: CCGGCCTAAGGTTAAGTCGCCCTCGCTCGAGCGAGGGCGACTTAACCTTAGGTTTTTG;

SCR-R: AATTCAAAAACCTAAGGTTAAGTCGCCCTCGCTCGAGCGAGGGCGACTTAACCTTAGG.

The target sequence for shATOH7-1 is GTGAAGTTACAGTATCCATTA available from TRC library database (TRCN0000423538) (Moffat et al., 2006), and the sequences for the hairpin pairs are as follows:

shATOH7-1F: CCGGGTGAAGTTACAGTATCCATTACTCGAGTAATGGATACTGTAACTTCACTTTTTTGAAT;

shATOH7-1R: AATTCAAAAAGTGAAGTTACAGTATCCATTACTCGAGTAATGGATACTGTAACTTCAC.

The target sequence for shATOH7-2 is GATTCTCAGATTACCTTTATT designed with an online tool (http://www.broad.mit.edu/genome_bio/trc/rnai.html) (Moffat et al., 2006), and the sequences of the hairpin pair are as follows:

shATOH7-2F:

CCGGGATTCTCAGATTACCTTTATTCTCGAGAATAAAGGTAATCTGAGAATCTTTTTG;

shATOH7-2R:

AATTCAAAAAGATTCTCAGATTACCTTTATTCTCGAGAATAAAGGTAATCTGAGAATC.

Both shRNAs are targeted to the 3′UTR of ATOH7 gene. The oligos of the hairpin pairs were purchased from Eurofins Genomics, annealed together, and cloned into AgeI and EcoRI cloning sites of pLKO.1-GFP-P2A-Puro or pLKO.1-RFP-P2A-Puro vector with Quick Ligation Kit (New England Biolabs, Cat#M2200S) (Moffat et al., 2006).

#### Production of shATOH7 lentivirus and virus validation

Lentivirus was produced with Lenti-X 293T packaging cells (Clontech, Cat#632180) transfected with viral vectors and packaging plasmids pMD2.G and psPAX2 (Addgene, Cat#12259 and Cat#12260) in DMEM medium (Wisent Bioproducts, Cat#319-005-CL) supplemented with 10% fetal bovine serum (FBS) (Wisent Bioproducts, Cat#080-150) (Sanjana et al., 2014). Briefly, for each virus, 5 × 10^6^ cells were seeded in a 10-cm plate the day before transfection. On the day of transfection, culture medium was removed and replaced with 5 ml of prewarmed fresh medium. A mixture of 3 μg viral vector, 2.25 μg pPAX2, 0.75 μg pMD2.G and 18 ul of Lipofectamine 2000 (Life Technologies, Cat#11668019) in 3.0 ml of Opti-MEM I Reduced-Serum Medium (1X) (Life Technologies, Cat#31985070) was added dropwise to the cells with gentle mixing by rocking the plate back and forth. The culture medium with transfection reagent was removed 12 hours later, and replaced with 10 ml of warm fresh medium. Virus-containing supernatants were harvested 48 hours after transfection, filtered through a 0.45 μm low protein binding membrane (Sarstedt, Cat#83.1826), and concentrated with Lenti-X Concentrator (Clontech, Cat#631231) according to the inserted protocol. Virus pellets were dissolved in 100 μl of IMDM (Wisent Bioproducts, Cat#319-105-CL), aliquoted and stored at −80°C.

For validation, a developing human cone cell line was generated by transduction of human retina with shRB1 virus and selection for replicating cells. Identity was validated with multiple cone markers, then cells were transduced with shATOH7 lentivirus for 8 days. To assess ATOH7 knockdown efficiency, RNA and protein were extracted from the cells, and Real-Time qRT-PCR as well as Western blot were carried out as described previously (Chen et al., 2009) to measure ATOH7 expression. Human β-ACTIN and TBP (TATA-Box Binding Protein) were used as qRT-PCR references, and β-ACTIN was used as a loading control for Western blot (Figure S7). The PCR primers are the following.

ATOH7-F: AGTACGAGACCCTGCAGATG
ATOH7-R: TGGAAGCCGAAGAGTCTCTG
β-ACTIN-F: AAAGCCACCCCACTTCTCTCTAA
β-ACTIN-R: ACCTCCCCTGTGTGGACTTG
TBP-F: ATGTTGAGTTGCAGGGTGTG
TBP-R: CAGCACGGTATGAGCAACTC

#### Transduction of developing human retinal explants with shATOH7 lentivirus

Developing human retinas at gestational age from GW17 to GW19 were sterilely dissected in IMDM and cut radially. Tissue fragments were transferred on to cell-culture inserts (Millpore, Cat#PICM03050) with photoreceptor side down (Jin and Xiang, 2012). Inserts with retinal fragments were quickly put in 6-well plates with 1300 μl of prewarmed retina culture medium, and incubated at 37°C with 5% CO_2_. Due to spatiotemporal difference in developing human retina development (Hoshino et al., 2017), each explant culture was co-transduced with both control and treatment viruses by adding freshly thawed virus mixture on top of retinal fragment (Figure 1A). The explants were cultured for about 3 weeks, and the medium was changed twice per week, with half of the culture medium replaced with fresh medium.

#### Immunofluorescence staining and microscopy of retinal explants

After about 3 weeks of culture, the retinal explants were fixed with 4% paraformaldehyde overnight at 4°C, washed 3 times with cold PBS, dehydrated in 30% sucrose for 24 hour at 4°C, embedded in 30% sucrose/OCT compound (1:1), and stored at −80°C. The frozen tissue blocks were sectioned at 16 μm thickness, and air-dried at room temperature for 4 hours, stored at −20°C. For immunostaining, sections were rinsed in Tris-buffered saline with 0.1% Tween 20 (TBS-T), incubated in blocking buffer containing 2.5% bovine serum albumin (BSA) in TBS-T for 1 hour at room temperature, and labeled with primary antibodies diluted in blocking buffer overnight at 4°C. The following day, sections were washed 3 times with TBS-T, and incubated for 1 hour at room temperature with appropriate secondary antibodies conjugated to 488/568/647 fluorophores. Subsequently, sections were washed with TBS-T for 3 times, cover-slipped, mounted in VECTASHIELD Hardset Antifade Mounting Medium with DAPI (Vector Laboratories, Cat# H-1500) to counterstain nuclei, and imaged using Nikon Ti-E inverted confocal microscope. Images were processed using ImageJ (NIH). For each explant per staining, at least 10 images were processed. Cones were confirmed with RXRγ or GNAT2 staining, and rods with NRL staining. The ratios of percentages of cones versus rods in the control and treatment virus transduced photoreceptors were calculated. Antibodies were used at the following dilutions: ATOH7 - 1:700, GFP - 1:300 (rabbit polyclonal; Life Technologies), GNAT2 - 1:100, mCherry - 1:500, NRL - 1:00, RFP 1:300, RXRγ - 1:100.

## Figure legends

**Figure S1 Related to Figure 1. Dataset metrics and immunohistochemistry of cell type markers.**

(A) Table showing the QC metrics (number of cells, mean UMI, mean number of genes), and tissue origin. (B-C) Distribution of (B) UMIs per cell and (C) Number of Genes Expressed in each sample. (D-E) UMAP embedding of entire human retinal dataset, including adult, and colored by (D) age and (E) annotated cell types. (F) Proportion of cell ages in each cell type. (G) Proportion of cell types at each age. (H-N) Immunohistochemistry of the developing human retina, detecting cell-type markers: (H) S-OPSIN (short wavelength cones); (I) L/M-OPSIN (long/medium wavelength cones); (J) RHO (rods); (K) PRKCA and VSX2 (bipolar cells); (L) BRN3A (RGCs) and CALRETININ (Horizontal, Amacrine and RGC cells.); and (M) Calbindin (Horizontal cells) and (N) Calbindin (Horizontal Cells) and Calretinin (Horizontal, Amacrine and RGC cells). Nuclei are counterstained with DAPI. Scale Bar: 50 μm. Abbreviations: Hgw - human gestational weeks; Hpnd - human postnatal day; RPCs - retinal progenitor cells; RGCs - retinal ganglion cells; AC/HC Pre - amacrine cell/horizontal cell precursors; BC/Photo Pre - bipolar cell/photoreceptor cell precursors; NBL - neuroblast layer; GCL - ganglion cell layer; ONL - outer nuclear layer; OPL - outer plexiform layer; INL - inner nuclear layer; IPL - inner plexiform layer.

**Figure S2 Related to Figure 2. Comparisons of temporal gene expression between human and mouse primary and neurogenic RPCs.**

(A-B) Density of pseudotime values of RPCs and Müller glia by (A) cell type and (B) age, with the dashed line demarking pseudotime threshold between Early and Late RPCs. (C) Proportion of RPCs in each cell cycle phase across human (left) and mouse (right) scRNA-Seq retinal development. (D) Immunostaining for MKI67 and RLBP1 co-localization in the central and peripheral retina across the developing human retina. Nuclei are counterstained with DAPI. Scale bar: 50μm. (E) Immunohistochemistry for SOX2 and RLBP1 in human retina across development. Nuclei are counterstained with DAPI. Scale bar: 50μm. (F) Bar chart showing the proportion of cells displaying co-localization of SOX2*+* and RLBP1*+* across development. (G-H) Heatmaps showing temporal gene expression within (G) RPCs and (H) Neurogenic RPCs. Highlighted genes are the human orthologs of genes highlighted in the the mouse scRNA-Seq primary and neurogenic RPC analyses. (I) Dotplot displaying the relative expression and proportions of cells expressing transcripts that display divergent temporal expression between mouse and human neurogenic cells. (J-K) Density plots of primary and neurogenic RPC pseudotime colored by (J) cell type or (K) developmental age. (L) Proportion of neurogenic cells in each cell cycle phase across human (left) and mouse (right) retinal scRNA-Seq datasets. Abbreviations: Hgw - human gestational weeks; GW - gestational weeks; Hpnd - human postnatal day; NBL - neuroblast layer; GCL - ganglion cell layer; ONL - outer nuclear layer; OPL - outer plexiform layer; INL - inner nuclear layer; IPL - inner plexiform layer; C - central retina; P - peripheral retina.

**Figure S3 Related to Figure 3. Pseudotime analysis of starburst amacrine cells and cone samples and cellular expression of pseudotime markers.**

(A-F) UMAP embedding and pseudotime heatmap for (A-C) amacrine and Starburst amacrine cells and (D-F) organoid cones (left branch) and *in vivo* cones (right branch) trajectories. (A-B; D-E) UMAP embeddings are colored by (A,D) pseudotime values, (B) cell type or (E) age. (C,F) In the pseudotime heatmap, cells are ordered by cell type and pseudotime and plotted genes are differentially expressed genes along pseudotime. (G) Heatmap showing cell type expression of all heatmap genes from Fig 3 and S3 in each cell type. Abbreviations: Hgw - human gestational weeks; Hpnd - human postnatal day; RPCs - retinal progenitor cells; AC/HC Pre - amacrine cell/horizontal cell precursors; Starburst/Starburst AC - starburst amacrine cells; BC/Photo Pre - bipolar cell/photoreceptor precursors, RGCs - retinal ganglion cells.

**Figure S4 Related to Figure 4. Identification of differential gene expression within retinal cell types across macula and peripheral cells.**

(A-H) Volcano plots (left) showing fold enrichment of genes between macula and peripheral regions, and heatmaps (right) of significantly enriched genes and their relative expression in the macular and peripheral samples ordered by region and developmental age. (I) Plots showing proportion of macula and peripheral cells of each cell type across (left) all regionally isolated cells, (middle) Hgw20 samples, and (right) Hpnd8 samples. (J-Q) Pseudotime heatmaps of the identified region-specific genes of each cell type and their correlation with pseudotime. Cells are ordered by age instead of pseudotime in (N). Abbreviations: Hgw - human gestational weeks; Hpnd - human postnatal day; RPCs - retinal progenitor cells; RGCs - retinal ganglion cells; AC/HC Pre - amacrine cell/horizontal cell precursors; BC/Photo Pre - bipolar cell/photoreceptor cell precursors.

**Figure S5 Related to Figure 5. Identification and validation of human horizontal cell subtype markers.**

(A-F) UMAP embedding of horizontal cells, colored by the relative gene expression of (A) *ONECUT1*, (B) *ONECUT2*, (C) *ONECUT3*, (D) *CALB1*, (E) *PCDH9*, and (F) cellular subtypes. (G) Immunostaining for LHX1 and ONECUT2 in the central (top) and peripheral (bottom) retina at Hgw9, 11 and 14, with magnified views of boxed regions. Scale bars: 150 μm in Hgw9 low magnification images; 50 μm; and 10 μm in highest magnification. (H) Proportion of ONECUT2+ cells that are also LHX1+ in central and peripheral retina at Hgw9, 11 and 14. (I) Heatmap showing cell type expression of highlighted marker genes of horizontal cell types and other cell types. Abbreviations: Commit HCs - committed horizontal cells; Hgw - human gestational weeks; Hpnd - human postnatal day; C - central retina; P - peripheral retina; NBL - neuroblast layer; GCL - ganglion cell layer; AC/HC Pre. - amacrine cell/horizontal cell precursors; RGCs - retinal ganglion cells; RPCs - retinal progenitor cells; BC/Photo Pre - bipolar cell/photoreceptor precursors.

**Figure S6. Related to Figure 6. Correlation heatmaps of scCoGAPS patterns to annotated cell features.**

Heatmaps displaying correlations of (A-D) human and (E-H) mouse patterns to (A,C,E,G) retinal cell types, (B,D,F,H) age and Total mRNAs in (A,B,E,F) human and (C,D,G,H) mouse datasets. Abbreviations: Hgw - human gestational weeks; Hpnd - human postnatal day; AC/HC Pre. - amacrine cell/horizontal cell precursors; RGCs - retinal ganglion cells; RPCs - retinal progenitor cells; BC/Photo Pre - bipolar cell/photoreceptor precursors; Photo Pre - photoreceptor precursors.

**Figure S7. Related to Figure 7. Spatiotemporal expression of ATOH7 in developing human retina.**

(A) Percentage of cells that express *ATOH7* in each cell type of the dataset. (B) Plot showing the distribution of *ATOH7* and *CRX* expression in each cell type. (C) ATOH7 and OTX2 immunostaining in central and peripheral human retina at Hgw16, 18 and 20, with magnified views of boxed regions. (D) ATOH7 and CRX immunostaining in central and peripheral human retina at Hgw14, Hgw18 Hgw20, and Hgw22, with magnified views of the boxed regions. (C-D) Scale bars: 50 μm and 10 μm (magnified views), except for the high magnification views in (C) Hgw16-C which represent 5 μm. Nuclei are counterstained with DAPI. (E) Relative *ATOH7* RNA expression in shATOH7 lentivirus transduced cells determined using RT-qPCR. (F) ATOH7 protein expression from virus transduced cells from (E). (G) Representative images from human retinal explants co-transduced with shSCR (GFP) and shATOH7-2 (RFP) lentiviruses and stained with cone marker RXRγ (blue). White arrows, RXRγ+ cells expressing shSCR; yellow arrows, RXRγ+ cells expressing shATOH7. Scale bars, 20μm. (H) Ratio of cones(RXRγ+)/rods(RXRγ-) in shATOH7 vs shSCR cells. Data are presented as means ± SD. (I) Representative images from human retinal explants co-transduced with shSCR (GFP) and shATOH7-2 (RFP) lentiviruses and stained with rod marker NRL (blue). White arrows, NRL+ cells expressing shSCR; yellow arrows, NRL+ cells expressing shATOH7. Scale bars, 20μm. (J) Ratio of cones(NRL-)/rods(NRL+) in shATOH7 vs shSCR cells. Data are presented as means ± SD. Abbreviations: AC/HC Pre - amacrine cell/horizontal cell precursors; BC/Photo Pre - bipolar cell/photoreceptor precursors; RGCs - retinal ganglion cells; RPCs - retinal progenitor cells; Precurs - precursors; Hgw - human gestational weeks; C - central retina; mid-P - mid-peripheral retina; P - peripheral retina.

**Table S1. Related to Figure 1.** Table of cell counts from scRNA-Seq experiments.

**Table S2. Related to Figures 1, S1, 3, S3, 5 and S5.** High variance genes used as input for dimension reductions and Scanpy analyses.

**Table S3. Related to Figures 3 and S3.** Input parameters for Scanpy analyses.

**Movie S1. Related to Figure 1B.** Movie of the 3D-UMAP dimension reduction embedding of the developing human retina scRNA-Seq dataset with cells colored by developmental age.

**Movie S2. Related to Figure 1C.** Movie of the 3D-UMAP dimension reduction embedding of the developing human retina scRNA-Seq dataset with cells colored by annotated cell type.

