## Supplemental Figures S1-S7 for "Single-cell analysis of human retina identifies evolutionarily conserved and species-specific mechanisms controlling development"

### Figure S1

A

| Organoids | # Cells | Mean UMI | Mean Genes | Sample Origin |
|---|---|---|---|---|
| **Day24** | 24 | 3454 | 1602 | JHMI |
| **Day30** | 175 | 5840 | 2145 | JHMI |
| **Day42** | 5919 | 3896 | 1682 | JHMI |
| **Day59** | 5424 | 3185 | 1331 | JHMI |

| Human Retina | # Cells | Mean UMI | Mean Genes | Sample Type | Sample Origin |
|---|---|---|---|---|---|
| **Hgw9** | 8974 | 4630 | 1701 | Whole Retina | Canada |
| **Hgw11** | 8058 | 4015 | 1683 | Whole Retina | Canada |
| **Hgw12** | 5467 | 2797 | 1040 | Whole Retina | China |
| **Hgw13** | 9912 | 3508 | 1459 | Whole Retina | Canada |
| **Hgw14** | 2958 | 3518 | 1507 | Whole Retina | China |
| **Hgw15** | 7480 | 2202 | 1063 | Whole Retina | Canada |
| **Hgw16** | 3196 | 5316 | 2091 | Whole Retina | China |
| **Hgw17** | 6034 | 2259 | 1132 | Whole Retina | Canada |
| **Hgw18** | 3565 | 4026 | 1720 | Whole Retina | China |
| **Hgw19_rep1** | 4332 | 2218 | 1082 | Whole Retina | Canada |
| **Hgw19_rep2** | 3899 | 2337 | 1176 | Whole Retina | Canada |
| **Hgw20_rep1** | 5060 | 4230 | 2005 | Macula | China |
| **Hgw20_rep2** | 7633 | 4180 | 1818 | Periphery | China |
| **Hgw22** | 2230 | 3557 | 1611 | Whole Retina | China |
| **Hgw24_rep1** | 2693 | 4224 | 1821 | Whole Retina | China |
| **Hgw24_rep2** | 4468 | 3832 | 1679 | Whole Retina | China |
| **Hgw27** | 2054 | 3442 | 1539 | Whole Retina | China |
| **Hpnd8_rep1** | 4010 | 3024 | 1413 | Macula | China |
| **Hpnd8_rep2** | 3372 | 2457 | 1261 | Periphery | China |
| **Adult (86yo)** | 11618 | 3350 | 1472 | Whole Retina | JHMI |

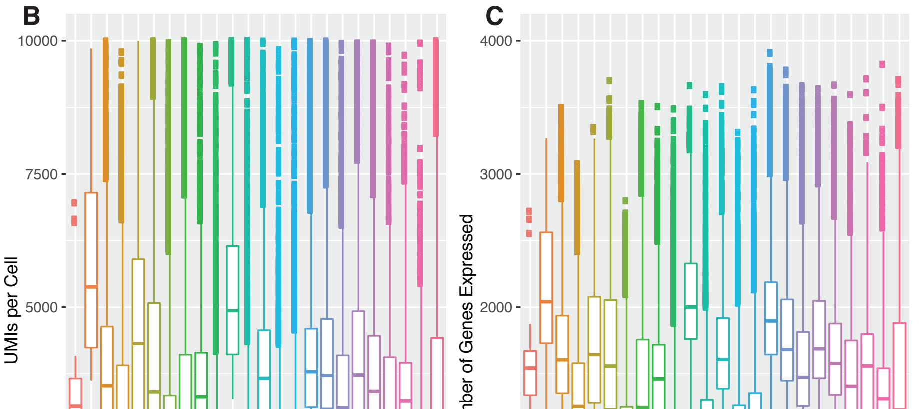


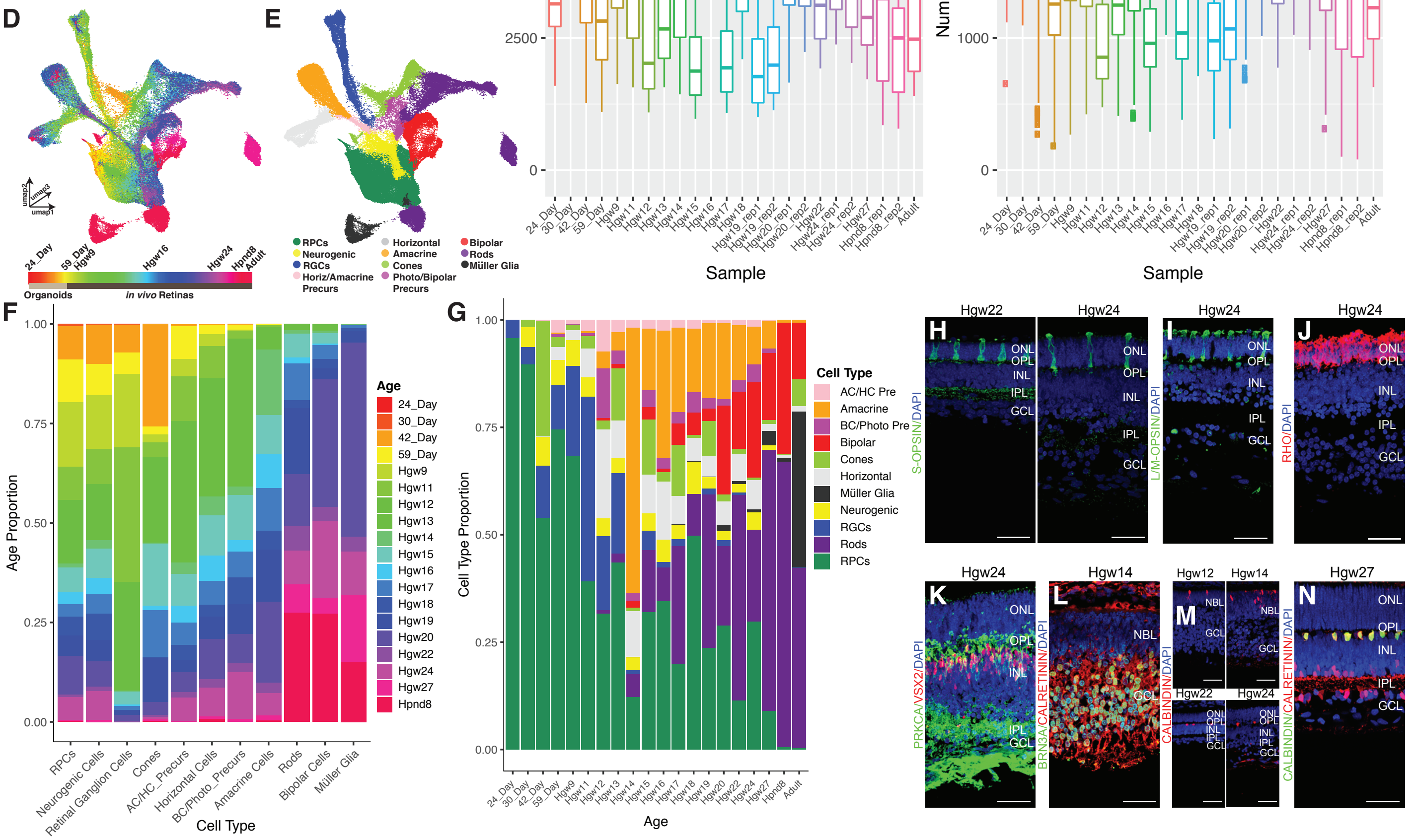


**Figure S1. Related to Figure 1. Dataset metrics and immunohistochemistry of cell type markers.** (A) Table showing the QC metrics (number of cells, mean UMI, mean number of genes), and tissue origin. (B-C) Distribution of (B) UMIs per cell and (C) Number of Genes Expressed in each sample. (D-E) UMAP embedding of entire human retinal dataset, including adult, and colored by (D) age and (E) annotated cell types. (F) Proportion of cell ages in each cell type. (G) Proportion of cell types at each age. (H-N) Immunohistochemistry of the developing human retina, detecting cell-type markers: (H) S-OPSIN (short wavelength cones); (I) L/M-OPSIN (long/medium wavelength cones); (J) RHO (rods); (K) PRKCA and VSX2 (bipolar cells); (L) BRN3A (RGCs) and CALRETININ (Horizontal, Amacrine and RGC cells.); and (M) Calbindin (Horizontal cells) and (N) Calbindin (Horizontal Cells) and Calretinin (Horizontal, Amacrine and RGC cells). Nuclei are counterstained with DAPI. Scale Bar: 50 µm. Abbreviations: Hgw - human gestational weeks; Hpnd - human postnatal day; RPCs - retinal progenitor cells; RGCs - retinal ganglion cells; AC/HC Pre - amacrine cell/horizontal cell precursors; BC/Photo Pre - bipolar cell/photoreceptor cell precursors; NBL - neuroblast layer; GCL - ganglion cell layer; ONL - outer nuclear layer; OPL - outer plexiform layer; INL - inner nuclear layer; IPL - inner plexiform layer.

### Figure S2

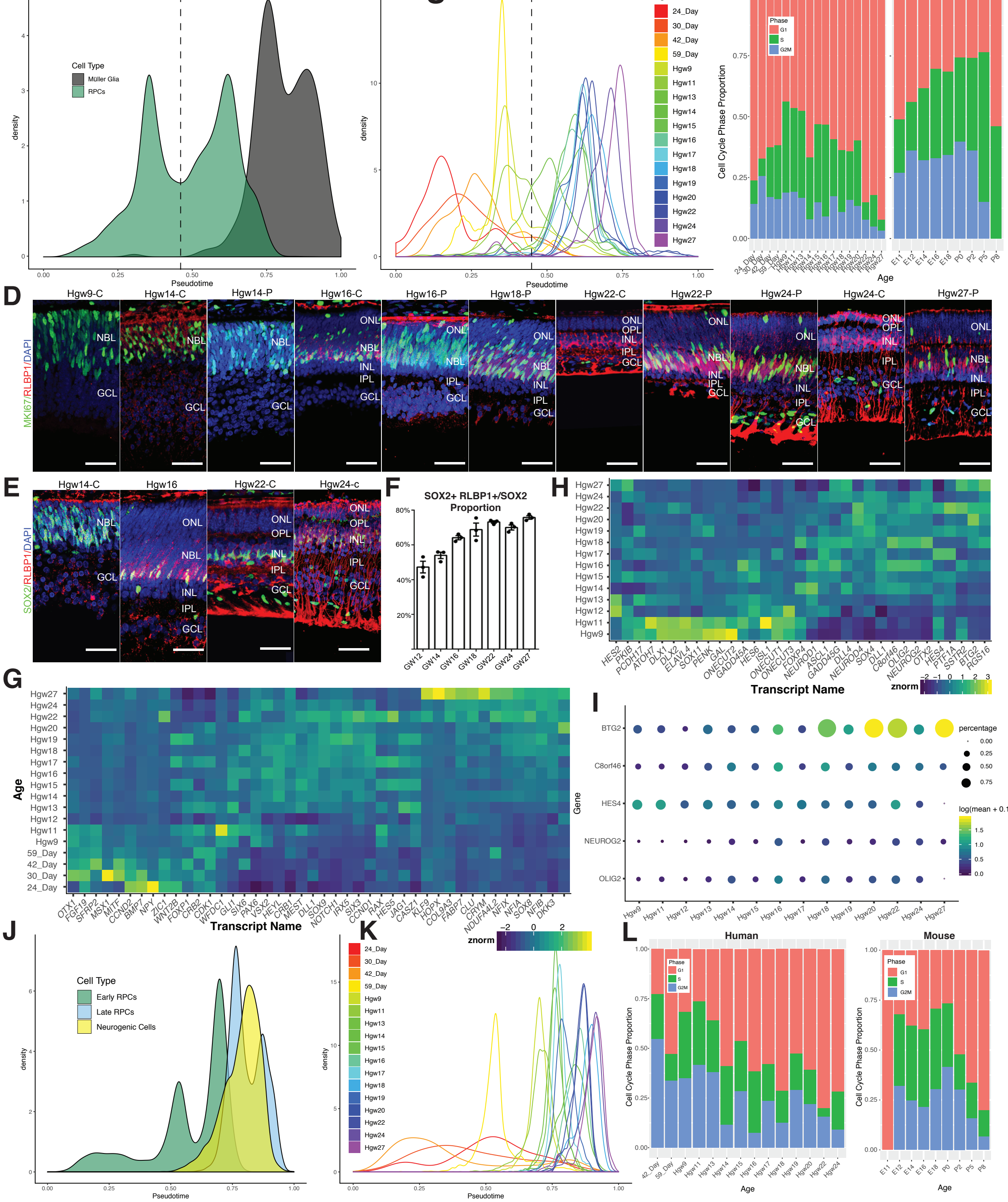


**Figure S2. Related to Figure 2. Comparisons of temporal gene expression between human and mouse primary and neurogenic RPCs.** (A-B) Density of pseudotime values of RPCs and Müller glia by (A) cell type and (B) age, with the dashed line demarking pseudotime threshold between Early and Late RPCs. (C) Proportion of RPCs in each cell cycle phase across human (left) and mouse (right) scRNA-Seq retinal development. (D) Immunostaining for MKI67 and RLBP1 co-localization in the central and peripheral retina across the developing human retina. Nuclei are counterstained with DAPI. Scale bar: 50µm. (E) Immunohistochemistry for SOX2 and RLBP1 in human retina across development. Nuclei are counterstained with DAPI. Scale bar: 50µm. (F) Bar chart showing the proportion of cells displaying co-localization of SOX2+ and RLBP1+ across development. (G-H) Heatmaps showing temporal gene expression within (G) RPCs and (H) Neurogenic RPCs. Highlighted genes are the human orthologs of genes highlighted in the the mouse scRNA-Seq primary and neurogenic RPC analyses. (I) Dotplot displaying the relative expression and proportions of cells expressing transcripts that display divergent temporal expression between mouse and human neurogenic cells. (J-K) Density plots of primary and neurogenic RPC pseudotime colored by (J) cell type or (K) developmental age. (L) Proportion of neurogenic cells in each cell cycle phase across human (left) and mouse (right) retinal scRNA-Seq datasets. Abbreviations: Hgw - human gestational weeks; GW - gestational weeks; Hpnd - human postnatal day; NBL - neuroblast layer; GCL - ganglion cell layer; ONL - outer nuclear layer; OPL - outer plexiform layer; INL - inner nuclear layer; IPL - inner plexiform layer; C - central retina; P - peripheral retina.

### Figure S3

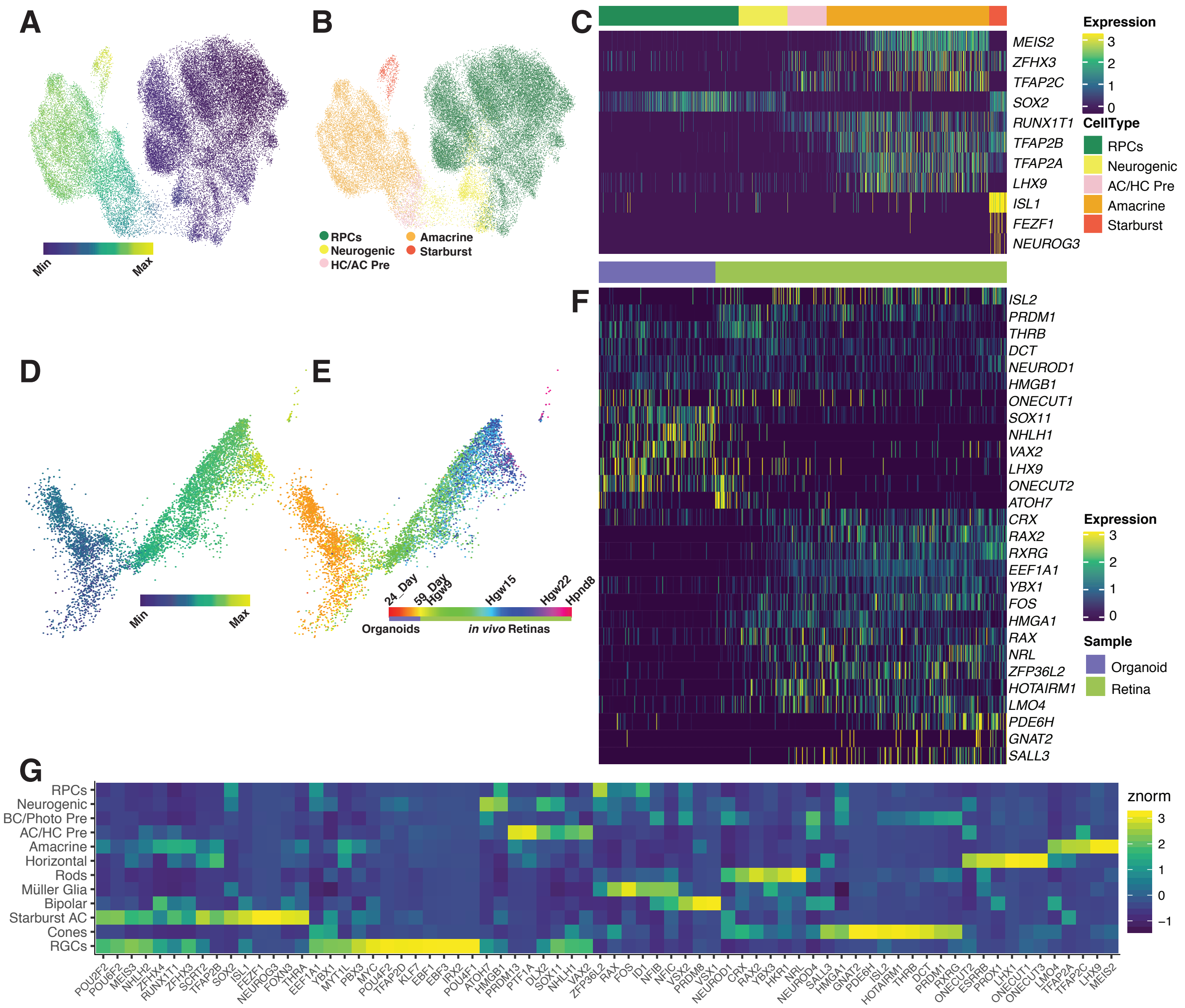


**Figure S3. Related to Figure 3. Pseudotime analysis of starburst amacrine cells and cone samples and cellular expression of pseudotime markers.** (A-F) UMAP embedding and pseudotime heatmap for (A-C) amacrine and Starburst amacrine cells and (D-F) organoid cones (left branch) and in vivo cones (right branch) trajectories. (A-B; D-E) UMAP embeddings are colored by (A,D) pseudotime values, (B) cell type or (E) age. (C,F) In the pseudotime heatmap, cells are ordered by cell type and pseudotime and plotted genes are differentially expressed genes along pseudotime. (G) Heatmap showing cell type expression of all heatmap genes from Fig 3 and S3 in each cell type. Abbreviations: Hgw - human gestational weeks; Hpnd - human postnatal day; RPCs - retinal progenitor cells; AC/HC Pre - amacrine cell/horizontal cell precursors; Starburst/ Starburst AC - starburst amacrine cells; BC/Photo Pre - bipolar cell/photoreceptor precursors, RGCs - retinal ganglion cells.

### Figure S4

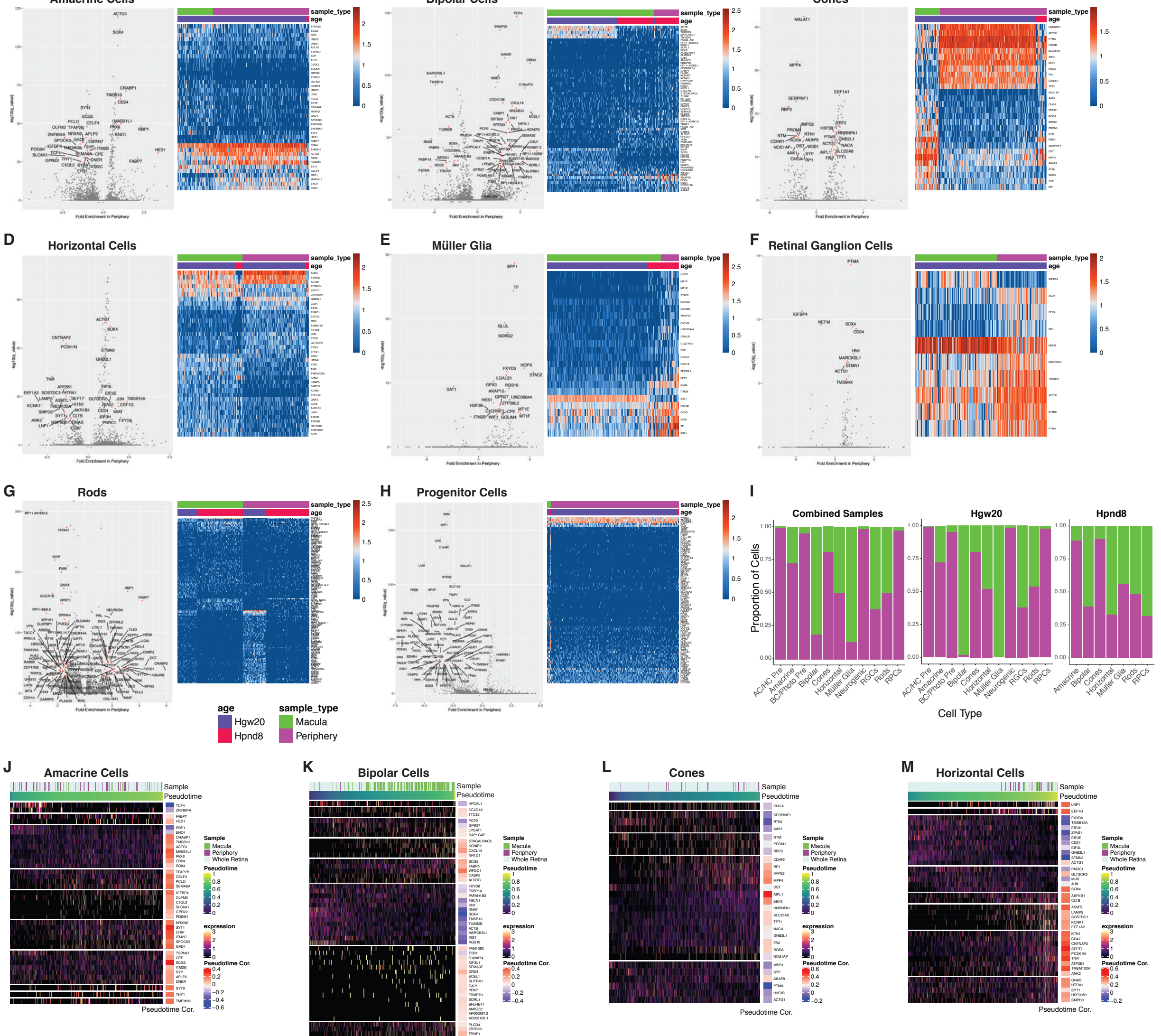


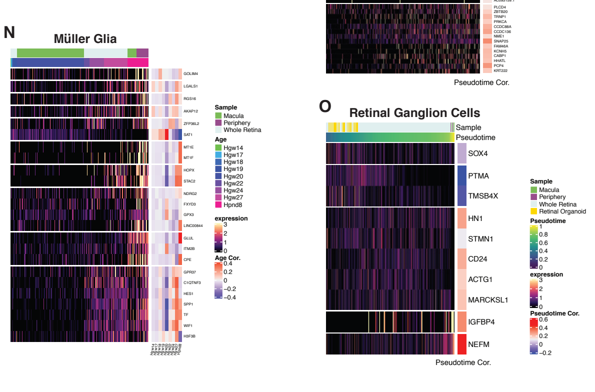


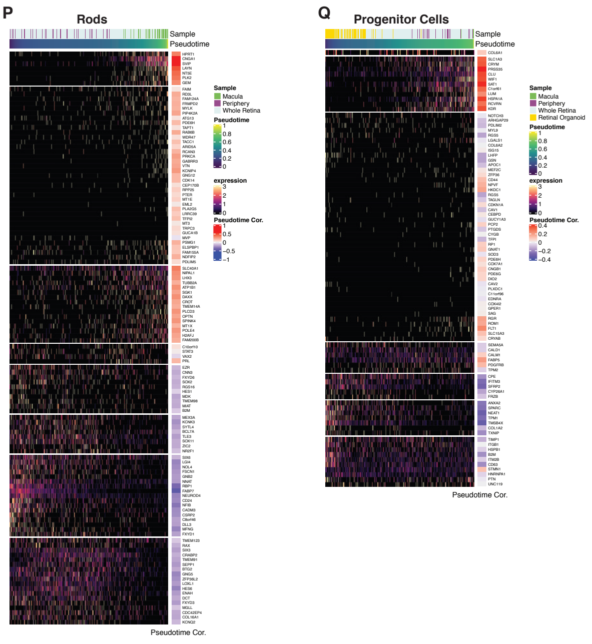


**(Next Page) Figure S4. Related to Figure 4. Identification of differential gene expression within retinal cell types across macula and peripheral cells.** (A-H) Volcano plots (left) showing fold enrichment of genes between macula and peripheral regions, and heatmaps (right) of significantly enriched genes and their relative expression in the macular and peripheral samples ordered by region and developmental age. (I) Plots showing proportion of macula and peripheral cells of each cell type across (left) all regionally isolated cells, (middle) Hgw20 samples, and (right) Hpnd8 samples. (J-Q) Pseudotime heatmaps of the identified region-specific genes of each cell type and their correlation with pseudotime. Cells are ordered by age instead of pseudotime in (N). Abbreviations: Hgw - human gestational weeks; Hpnd - human postnatal day; RPCs - retinal progenitor cells; RGCs - retinal ganglion cells; AC/HC Pre - amacrine cell/horizontal cell precursors; BC/Photo Pre - bipolar cell/photoreceptor cell precursors.

### Figure S5

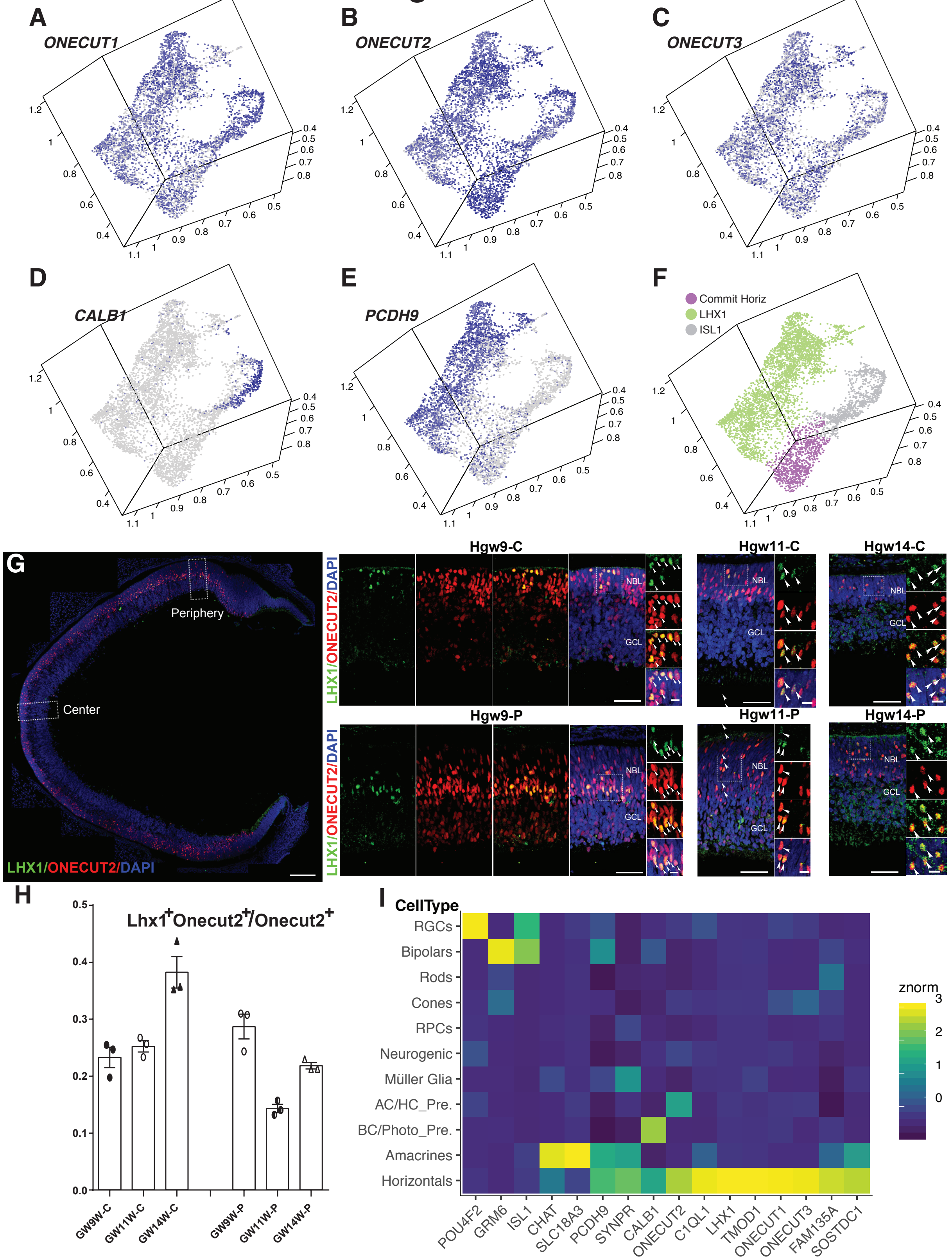


**Figure S5. Related to Figure 5. Identification and validation of human horizontal cell subtype markers.** (A-F) UMAP embedding of horizontal cells, colored by the relative gene expression of (A) ONECUT1, (B) ONECUT2, (C) ONECUT3, (D) CALB1, (E) PCDH9, and (F) cellular subtypes. (G) Immunostaining for LHX1 and ONECUT2 in the central (top) and peripheral (bottom) retina at Hgw9, 11 and 14, with magnified views of boxed regions. Scale bars: 150 μm in Hgw9 low magnification images; 50 μm; and 10 μm in highest magnification. (H) Proportion of ONECUT2+ cells that are also LHX1+ in central and peripheral retina at Hgw9, 11 and 14. (I) Heatmap showing cell type expression of highlighted marker genes of horizontal cell types and other cell types. Abbreviations: Commit HCs - committed horizontal cells; Hgw - human gestational weeks; Hpnd - human postnatal day; C - central retina; P - peripheral retina; NBL - neuroblast layer; GCL - ganglion cell layer; AC/HC Pre. - amacrine cell/horizontal cell precursors; RGCs - retinal ganglion cells; RPCs - retinal progenitor cells; BC/Photo Pre - bipolar cell/photoreceptor precursors.

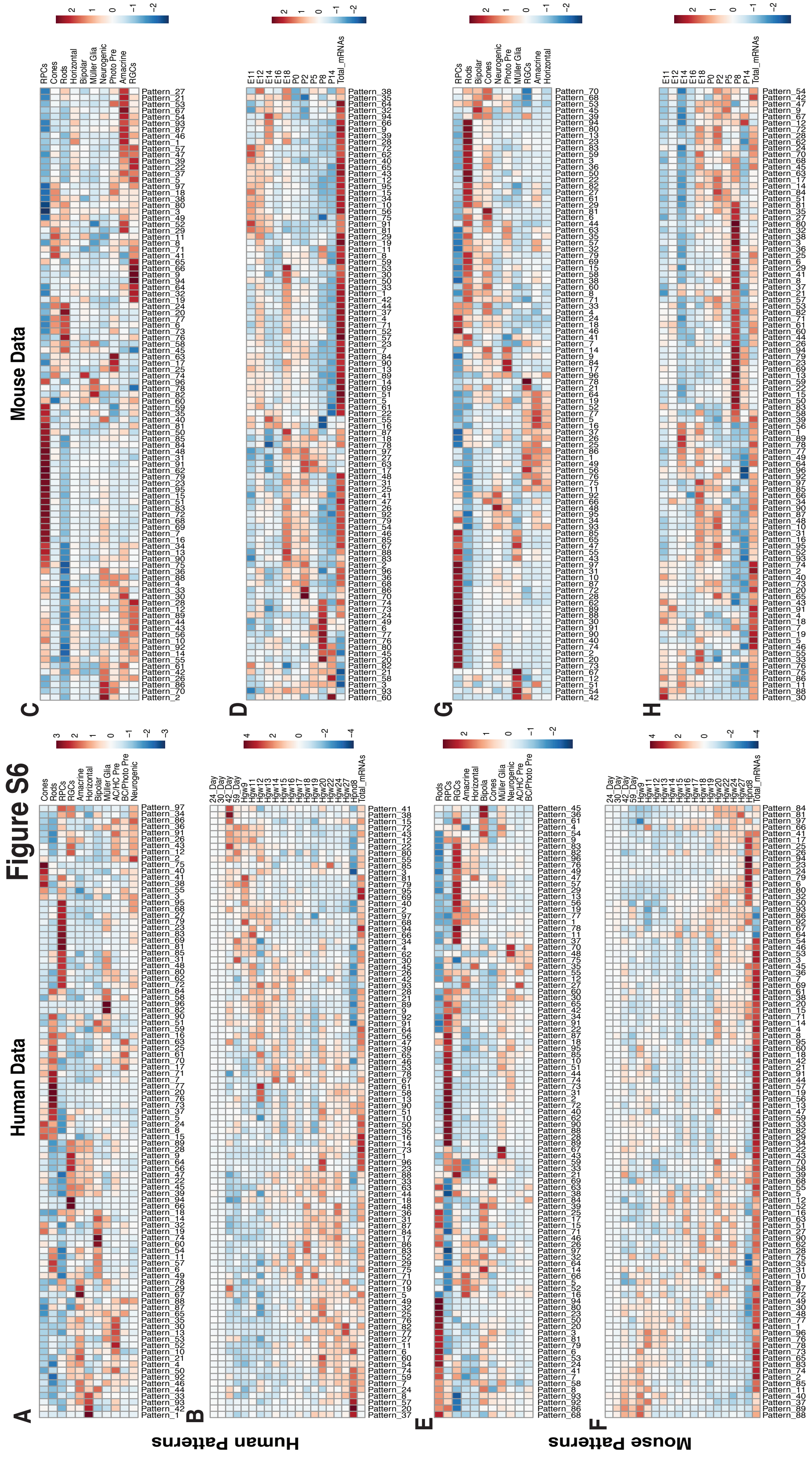


**Figure S6. Related to Figure 6. Correlation heatmaps of scCoGAPS patterns to annotated cell features.** Heatmaps displaying correlations of (A-D) human and (E-H) mouse patterns to (A,C,E,G) retinal cell types, (B,D,F,H) age and Total mRNAs in (A,B,E,F) human and (C,D,G,H) mouse datasets. Abbreviations: Hgw - human gestational weeks; Hpnd - human postnatal day; AC/HC Pre. - amacrine cell/horizontal cell precursors; RGCs - retinal ganglion cells; RPCs - retinal progenitor cells; BC/Photo Pre - bipolar cell/photoreceptor precursors; Photo Pre - photoreceptor precursors.

### Figure S7

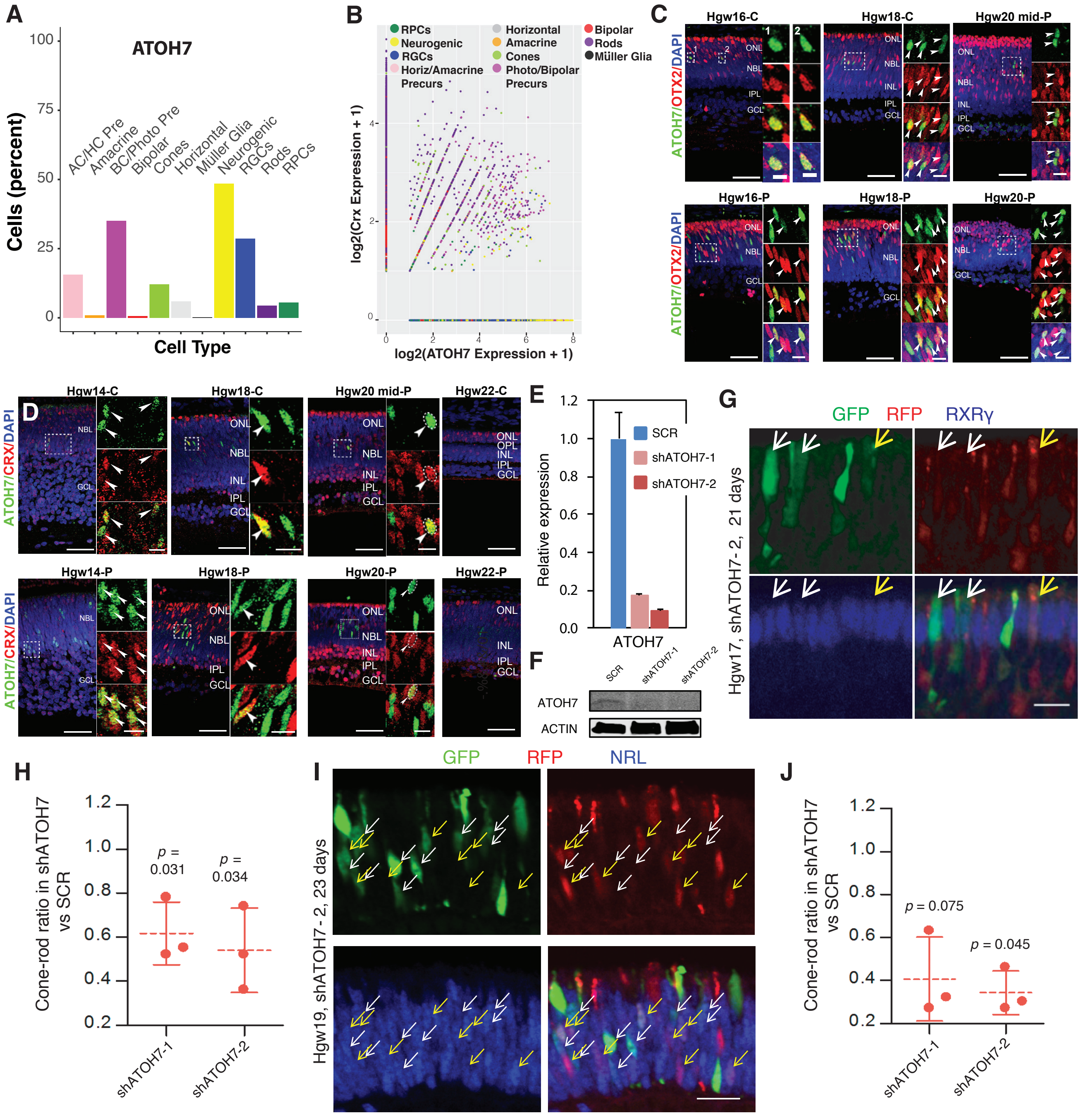


**Figure S7. Related to Figure 7. Spatiotemporal expression of ATOH7 in developing human retina.** A) Percentage of cells that express ATOH7 in each cell type of the dataset. (B) Plot showing the distribution of ATOH7 and CRX expression in each cell type. (C) ATOH7 and OTX2 immunostaining in central and peripheral human retina at Hgw16, 18 and 20, with magnified views of boxed regions. (D) ATOH7 and CRX immunostaining in central and peripheral human retina at Hgw14, Hgw18 Hgw20, and Hgw22, with magnified views of the boxed regions. (C-D) Scale bars: 50 μm and 10 μm (magnified views), except for the high magnification views in (C) Hgw16-C which represent 5 μm. Nuclei are counterstained with DAPI. (E) Relative ATOH7 RNA expression in shATOH7 lentivirus transduced cells determined using RT-qPCR. (F) ATOH7 protein expression from virus transduced cells from (E). (G) Representative images from human retinal explants co-transduced with shSCR (GFP) and shATOH7-2 (RFP) lentiviruses and stained with cone marker RXRγ (blue). White arrows, RXRγ+ cells expressing shSCR; yellow arrows, RXRγ+ cells expressing shATOH7. Scale bars, 20μm. (H) Ratio of cones(RXRγ+)/rods(RXRγ-) in shATOH7 vs shSCR cells. Data are presented as means ± SD. (I) Representative images from human retinal explants co-transduced with shSCR (GFP) and shATOH7-2 (RFP) lentiviruses and stained with rod marker NRL (blue). White arrows, NRL+ cells expressing shSCR; yellow arrows, NRL+ cells expressing shATOH7. Scale bars, 20μm. (J) Ratio of cones(NRL-)/rods(NRL+) in shATOH7 vs shSCR cells. Data are presented as means ± SD. Abbreviations: AC/HC Pre - amacrine cell/horizontal cell precursors; BC/Photo Pre - bipolar cell/photoreceptor precursors; RGCs - retinal ganglion cells; RPCs - retinal progenitor cells; Precurs - precursors; Hgw - human gestational weeks; C - central retina; mid-P - mid-peripheral retina; P - peripheral retina.
