## Supplemental Table S1 for "Single-cell analysis of human retina identifies evolutionarily conserved and species-specific mechanisms controlling development"

**Table S1 - Table of cell counts from scRNA-seq experiments**

| | | | | Cells Removed | | | | | | | | | | | | | | |
|---|---|---|---|---|---|---|---|---|---|---|---|---|---|---|---|---|---|---|
| **Sample** | **Sample type** | **Total Cells** | **Non-Retina** | **>=10,000 Transcripts** | **Doublets** | **FINAL CELL NUMBER** | | **RPCs** | **Neurogenic** | **RGCs** | **AC/HC Pre** | **Horizontal** | **Cones** | **Amacrine** | **Photo/BC Pre** | **Rods** | **Bipolar** | **Muller Glia** |
| 24_Day | Organoid | 9871 | 9847 | 0 | 0 | 24 | 24 | 23 | 0 | 1 | 0 | 0 | 0 | 0 | 0 | 0 | 0 | 0 |
| 30_Day | Organoid | 2255 | 1973 | 107 | 0 | 175 | 175 | 157 | 8 | 7 | 0 | 0 | 3 | 0 | 0 | 0 | 0 | 0 |
| 42_Day | Organoid | 7735 | 1732 | 84 | 0 | 5919 | 5919 | 3194 | 397 | 719 | 9 | 7 | 1588 | 1 | 4 | 0 | 0 | 0 |
| 59_Day | Organoid | 5612 | 151 | 37 | 0 | 5424 | 5424 | 4045 | 315 | 559 | 157 | 165 | 125 | 25 | 32 | 0 | 1 | 0 |
| Hgw9 | Whole Retina | 9428 | 43 | 408 | 3 | 8974 | 8974 | 6129 | 547 | 1885 | 84 | 201 | 117 | 5 | 6 | 0 | 0 | 0 |
| Hgw11 | Whole Retina | 8396 | 37 | 300 | 1 | 8058 | 8058 | 3153 | 354 | 3470 | 213 | 545 | 236 | 30 | 51 | 3 | 3 | 0 |
| Hgw12 | Whole Retina | 5526 | 7 | 45 | 7 | 5467 | 5467 | 1731 | 222 | 942 | 398 | 1137 | 111 | 218 | 632 | 45 | 31 | 0 |
| Hgw13 | Whole Retina | 10525 | 10 | 471 | 132 | 9912 | 9912 | 4315 | 348 | 1850 | 278 | 863 | 1212 | 421 | 317 | 211 | 97 | 0 |
| Hgw14 | Whole Retina | 3123 | 14 | 78 | 73 | 2958 | 2958 | 361 | 90 | 26 | 54 | 316 | 22 | 1825 | 52 | 160 | 51 | 1 |
| Hgw15 | Whole Retina | 7608 | 12 | 23 | 93 | 7480 | 7480 | 2397 | 294 | 341 | 152 | 677 | 961 | 1071 | 290 | 1079 | 218 | 0 |
| Hgw16 | Whole Retina | 3805 | 6 | 531 | 72 | 3196 | 3196 | 1104 | 166 | 38 | 79 | 432 | 72 | 950 | 75 | 253 | 27 | 0 |
| Hgw17 | Whole Retina | 6181 | 13 | 39 | 95 | 6034 | 6034 | 1201 | 198 | 99 | 108 | 404 | 720 | 1182 | 164 | 1660 | 295 | 3 |
| Hgw18 | Whole Retina | 3817 | 7 | 154 | 91 | 3565 | 3565 | 1777 | 266 | 1 | 73 | 197 | 22 | 525 | 169 | 344 | 188 | 3 |
| Hgw19_rep1 | Whole Retina | 4485 | 16 | 33 | 104 | 4332 | 4332 | 1022 | 119 | 65 | 20 | 187 | 371 | 762 | 94 | 1541 | 144 | 7 |
| Hgw19_rep2 | Whole Retina | 4086 | 6 | 56 | 125 | 3899 | 3899 | 925 | 98 | 51 | 38 | 198 | 310 | 682 | 100 | 1402 | 90 | 5 |
| Hgw20_rep1 | Macula | 5581 | 6 | 314 | 201 | 5060 | 5060 | 78 | 5 | 110 | 1 | 327 | 41 | 621 | 9 | 1083 | 2588 | 197 |
| Hgw20_rep2 | Periphery | 8424 | 5 | 432 | 354 | 7633 | 7633 | 3592 | 253 | 67 | 92 | 353 | 164 | 1622 | 177 | 1266 | 47 | 0 |
| Hgw22 | Whole Retina | 2454 | 10 | 144 | 70 | 2230 | 2230 | 255 | 44 | 14 | 28 | 139 | 29 | 284 | 60 | 1066 | 296 | 15 |
| Hgw24_rep1 | Whole Retina | 2984 | 2 | 199 | 90 | 2693 | 2693 | 782 | 111 | 0 | 52 | 179 | 25 | 195 | 131 | 708 | 503 | 7 |
| Hgw24_rep2 | Whole Retina | 4888 | 18 | 238 | 164 | 4468 | 4468 | 1353 | 184 | 0 | 60 | 311 | 18 | 434 | 166 | 821 | 1084 | 37 |
| Hgw27 | Whole Retina | 2232 | 14 | 108 | 56 | 2054 | 2054 | 185 | 21 | 0 | 4 | 35 | 18 | 132 | 21 | 1249 | 321 | 68 |
| Hpnd8_rep1 | Macula | 4167 | 138 | 16 | 3 | 4010 | 4010 | 28 | 0 | 0 | 0 | 39 | 2 | 5 | 0 | 2536 | 1373 | 27 |
| Hpnd8_rep2 | Periphery | 3424 | 17 | 18 | 17 | 3372 | 3372 | 13 | 0 | 0 | 0 | 19 | 18 | 41 | 0 | 2371 | 876 | 34 |
| Adult | Whole Retina | 12065 | 82 | 365 | 0 | 11618 | 11618 | 37 | 0 | 16 | 0 | 151 | 720 | 74 | 0 | 4879 | 1528 | 4213 |
| | | 113199 | | | | 107013 | | | | | | | | | | | | |
| | | | | | | 11542 | | | | | | | | | | | | |
| | **Total Cell Number** | **138,672** | **14,166** | **4,200** | **1,751** | **118,555** | **118,555** | **37,857** | **4,040** | **10,261** | **1,900** | **6,882** | **6,905** | **11,105** | **2,550** | **22,677** | **9,761** | **4,617** |

| | | |
|---|---|---|
| | **Final Retinal Cells** | 118,555 |
