## Supplemental Table S3 for "Single-cell analysis of human retina identifies evolutionarily conserved and species-specific mechanisms controlling development"

**Table S3 - Input parameters for Scanpy**

| Trajectory | Batch Correction | Scanpy Variable Genes Parameters | Neighbors for Diffusion Map | Number of Diffusion Components for Diffusion Pseudotime |
|---|---|---|---|---|
| Amacrine Cells | None | Seurat default parameters | UMAP (30 canberra) | 15 |
| Bipolar vs. Photoreceptors | Combat (Pedersen et al. 2012) between FACS and non-FACS sorted cells, scanpy regress_out Total_mRNAs | Cell Ranger, 3000 top genes | UMAP (100 euclidean) | 8 |
| Horizontal Cells | BBKNN between FACS and non-FACS sorted cells. | Seurat, default parameters | BBKNN (neighbors within batch = 30, trim=0, metric = angular) | 15 |
| Cones vs. Rods | Same as Horizontal Cells | Seurat, max_mean=4, n_bins=14 | BBKNN (default parameters, trim= 100) | 7 |
| Retinal Ganglion Cells | None | Cell Ranger, 2000 top genes | UMAP (50 canberra) | 15 |
| Organoid vs. Human Retina Cones | None | Seurat, default | UMAP (7 canberra) | 15 |
| Muller Glia | BBKNN between organoids, FACS and non-FACS sorted cells. | None | BBKNN (neighbors within batch = 10, trim = 0, n_trees=20) | 15 |
| Neurogenic Cells | Monocle 3 preprocessCDS between organoids, FACS and non-FACS sorted cells | Seurat, default parameter | UMAP(15 canberra) | 15 |
